# B cell humoral response and differentiation is regulated by the non-canonical poly(A) polymerase TENT5C

**DOI:** 10.1101/686683

**Authors:** Aleksandra Bilska, Monika Kusio-Kobiałka, Paweł S. Krawczyk, Olga Gewartowska, Bartosz Tarkowski, Kamil Kobyłecki, Jakub Gruchota, Ewa Borsuk, Andrzej Dziembowski, Seweryn Mroczek

**Author notes:** These authors contributed equally (Aleksandra Bilska, Monika Kusio-Kobiałka). Correspondence (A.D.), (S.M.).

## Abstract

TENT5C is a non-canonical cytoplasmic poly(A) polymerase (ncPAP) upregulated in activated B cells and suppressing their proliferation. Herein we measured the global distribution of poly(A) tail lengths in responsive B cells using a modified Nanopore direct RNA-sequencing approach and revealed that TENT5C polyadenylates immunoglobulin mRNAs regulating their steady-state levels. Consequently, TENT5C deficient B cells secrete less antibodies and KO mice have diminished gamma globulin concentrations despite the increased number of CD138^high^ plasma cells as a consequence of accelerated differentiation. TENT5C is explicitly upregulated in differentiating plasma cells by innate signaling. Importantly, TENT5C deficiency in B lymphocytes impairs the capacity of the secretory pathway through the reduction of ER volume and downregulation of unfolded protein response.

Our findings define the role of the TENT5C enzyme in B cell physiology and discover the first ncPAP engaged in the regulation of immunoglobulin mRNA poly(A) tails, thus serving as a regulator of humoral immunity.

## Introduction

Development of an adaptive humoral immune response requires activation of resting B cells following antigen recognition. This process is associated with structural and functional changes leading to the generation of high-affinity memory B cells and antibody-secreting plasma cells (ASC). Extensive B cell differentiation is characterized by the clonal expansion, somatic hypermutation leading to affinity maturation, isotype switching, and formation of ASC or memory cells. At a cellular level, this process involves the reorganization of the rough endoplasmic reticulum (ER) and Golgi compartments to promote immunoglobulin synthesis, assembly and secretion (Lynes and Simmen, 2011; Wiest et al., 1990). These global physiological changes occurring during B cell differentiation and activation are linked to broad changes in the transcriptomic profile which is controlled by the coordinated action of regulatory networks of transcriptional factors such as NF-kB, BCL6, IRF4, and BLIMP1 (De Silva and Klein, 2015). Recent studies have also revealed the involvement of RNA-binding proteins (RBPs) and microRNAs (miRNAs) in shaping the B cell transcriptome (Danger et al., 2014; Diaz-Munoz et al., 2017), suggesting that post-transcriptional gene expression regulation plays an important role in B cell physiology (Koralov et al., 2008; Thai et al., 2007; Vigorito et al., 2007; Xu et al., 2012). In addition to global transcript changes, B cell maturation is associated with a large increase in the translation of mRNAs targeted to the ER (Goldfinger et al., 2011; Wiest et al., 1990).

Essentially, every mRNA molecule, except histone mRNAs, is polyadenylated during 3’ end processing. Nuclear polyadenylation is mediated by canonical poly(A) polymerases that interact with 3’ end cleavage machinery. The poly(A) tail plays a critical role in mRNA stability and translation efficacy as nearly all mRNA decay pathways begin with the removal of poly(A) tails (Houseley and Tollervey, 2009; Hrit et al., 2014; Siwaszek et al., 2014). Previous studies of polyadenylation in B cells have largely focused on the role of alternative polyadenylation and splicing in the regulation of immunoglobulin isoforms (Enders et al., 2014; Peng et al., 2017; Pioli et al., 2014; Takagaki and Manley, 1998). However, in addition to the polyadenylation that occurs in the nucleus, the poly(A) tail can be expanded in the cytoplasm by non-canonical poly(A) polymerases (ncPAPs). This process is considered to play an important role in the activation of dormant deadenylated mRNAs during gametogenesis (Friday and Keiper, 2015) and in neuronal processes but has not yet been studied in B cells. We and others have recently identified a novel metazoan-specific family of cytoplasmic poly(A) polymerases, TENT5 (previously known as FAM46). In mammals, this family has 4 members (Kuchta et al., 2009; Kuchta et al., 2016; Mroczek et al., 2017), among which TENT5C is the best-characterized. The importance of TENT5C is underscored by the occurrence of TENT5C somatic mutations in about 20% of cases of multiple myeloma (MM) patients. Further work revealed that TENT5C is a *bona fide* MM cell growth suppressor (Mroczek et al., 2017; Zhu et al., 2017). TENT5C polyadenylates multiple mRNAs with a strong specificity to those encoding ER-targeted proteins. This partially explains TENT5C toxicity to MM cells since an increased protein load caused by the stabilization of ER-targeted mRNAs enhances the ER stress, to which MM is very sensitive. Initial characterization of TENT5C KO in mice revealed that it might play a role in the physiology of normal B cells since isolated primary splenocytes from TENT5C KO mice proliferate faster upon activation than those isolated from WT animals (Mroczek et al., 2017).

Here we studied the role of TENT5C in B cells in more detail. Using Nanopore direct RNA sequencing we have carried out a global analysis of poly(A) tail distribution in B cells from WT and TENT5C KO animals and determined that the primary targets of TENT5C are mRNAs encoding immunoglobulins (Ig). Basically, mRNAs encoding all classes of Ig had shorter poly(A) tails. The analysis also allowed us to draw some general conclusions about poly(A) tail dynamics such as the positive correlation between the length of the poly(A) tail and the mRNA expression level, which was previously questioned based on methods employing Illumina sequencing (Lima et al., 2017).

Importantly, further studies revealed that the production of immunoglobulins is lower in TENT5C KO B cells, leading to decreased gamma globulin concentrations in KO mice serum. TENT5C deficient cells are characterized by accelerated growth rate and faster differentiation to CD138^high^ plasma cells, which explains their increased number in the bone marrow (BM) and spleen of KO mice. Accordingly, TENT5C expression is limited to late stages of B cell lineage differentiation and is highly upregulated by innate signaling via specific Toll-like receptors (TLR). Despite the acceleration of B cell proliferation rate, a lack of TENT5C resulted in a decrease of ER compartment volume, reduced dynamic of its expansion during B cell activation, and downregulation of unfolded protein response. This, together with a decreased steady-state level of IgG mRNAs explain why TENT5C KO cells produce and secrete less antibodies.

In aggregate, we revealed that cytoplasmic polyadenylation by ncPAP TENT5C regulates the humoral immune response.

## Results

### Direct RNA sequencing reveals IgG mRNAs as specific TENT5C targets

TENT5C is implicated in the polyadenylation of mRNAs encoding proteins passing through the ER in multiple myeloma (MM) cells, which originate from terminally differentiated B cells (Mroczek et al., 2017). In order to identify TENT5C substrates in activated B cells, we implemented Oxford Nanopore Technologies (ONT) direct full-length RNA sequencing to measure poly(A) tail length at a genome-wide scale. Unlike traditional RNA-seq techniques, the Nanopore-based system detects DNA or RNA single molecules as they traverse through protein channels, without the need for an enzymatic synthesis reaction. Moreover, this sequencing strategy avoids limitations and biases introduced during amplification of long homopolymers such as adenine tracts within poly(A) tails as PCR amplification of cDNA is not required during library preparation (Feng et al., 2015; Garalde et al., 2018) (Figure 1A). In the case of RNA sequencing, the substrate is a whole single RNA molecule (or an RNA-DNA hybrid after optional reverse transcription) with the motor protein attached to its 3’-end, which in an ATP-dependent manner passes the RNA strand through the pore at a consistent rate. As the sequencing proceeds in the 3’ to 5’ direction, the adaptor oligo is detected first, followed by the poly(A) tail, then the entire body of the transcript is sequenced. For efficient sequencing, pure mRNA fractions are needed, and to avoid any biases total RNA was subjected to an mRNA enrichment-step using the mutated recombinant elongation initiation factor 4E (GST-eIF4E^K119A^) which has a high affinity to 5’-cap structure (Bajak and Hagedorn, 2008; Choi and Hagedorn, 2003) (Supplementary Figure 1A-B). According to our experience and previous reports, this is the most effective strategy for mRNA enrichment (Choi and Hagedorn, 2003), as the efficiency of mRNA enrichment and depletion of other unwanted high-abundance RNA species was estimated by qPCR and northern blot analysis for selected transcripts (Supplementary Figure 1C-E). Next, samples depleted of most of small non-coding RNAs and rRNAs, were subjected to one round of ribodepletion (Supplementary Figure 1C). RNA prepared in such a way isolated from LPS/IL4 activated, spleen-derived B cells (WT and TENT5C KO mice) was adapted for Nanopore direct-RNA sequencing with the MinION device (Garalde et al., 2018) (Figure 1A).

**Figure 1.**
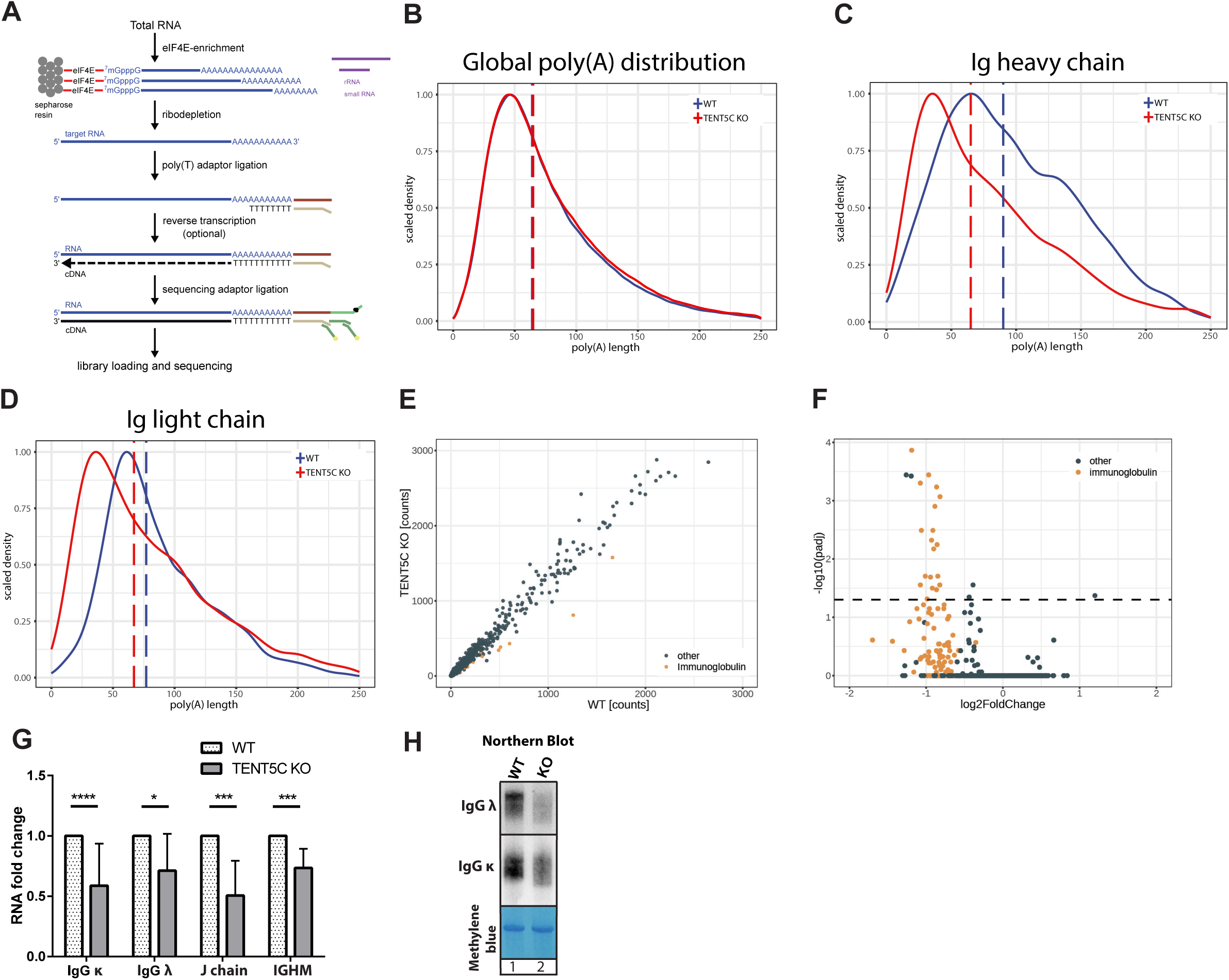
TENT5C regulates the steady-state level of immunoglobulins mRNAs through polyadenylation. (A) Library preparation workflow for Nanopore direct RNA sequencing. (B) (C) (D) Nanopore-based poly(A) lengths profiling of B cells isolated from WT and TENT5C KO, activated with LPS and IL4 for 7 days. Density distribution plots for all transcripts (B), Ig heavy chains transcripts (C) and Ig light chains (kappa) transcripts (D) scaled to a maximum of 1, are shown. Vertical dashed lines represent median poly(A) lengths for each condition. (E) Scatter plot of raw counts for individual transcripts obtained with Nanopore direct RNA sequencing for WT (x-axis) and TENT5C KO (y-axis). Immunoglobulins transcripts are marked in orange. (F) Volcano plot showing the results of DESeq2 differential expression analysis on data from Illumina sequencing of RNA isolated from B cells of WT and TENT5C KO animals activated with LPS and IL4 for 7 days. Immunoglobulins transcripts are marked in orange, dashed line marks the P value significance threshold (0.05). (G) Reverse transcription-qPCR analysis of immunoglobulins mRNA expression in WT and TENT5C KO cells. Values are shown as fold changes to WT. P values were calculated using unpaired Student’s t-test (n≥6). (H) Northern Blot analysis of κ and λ IgG mRNAs isolated from WT (lane 1) and TENT5C KO (lane 2) B cells activated with LPS and IL4 for 7 days.

Using ONT sequencing, we have generated 1.5M of transcriptome-wide full-length native-strand mRNA reads, which provided reliable information about steady-state poly(A) tail length in responsive B cells. We observed no global change in the mRNA polyadenylation status between WT and TENT5C deficient cells (Figure 1B). However, mRNAs encoding immunoglobulins had their median length of the poly(A) tails significantly decreased from 83 in WT to 67 adenosines in KO cells. This was observed for transcripts encoding both heavy and light immunoglobulin chains (Figure 1C, D). The effect was highly specific to IgG transcripts as poly(A) tails of other highly abundant mRNAs such as ribosomal proteins or mitochondrial transcripts were not affected at all (Supplementary Figure 1F, G). Finally, differential expression analysis based on ONT data showed that the abundance of Ig mRNAs was also decreased in TENT5C KO cells (Figure 1E, Supplementary Dataset 1).

In parallel with direct RNA-seq, we performed standard Illumina RNA sequencing, allowing us to perform a comparative correlation analysis between datasets. Again, Ig mRNAs were the most downregulated ones in TENT5C KO (Figure 1F, Supplementary Dataset 2), which was additionally confirmed by RT-qPCR (Figure 1G). Such strong specificity of TENT5C for IgG mRNAs is also clearly visible on the correlation scatterplot between the poly(A) tail length change and expression fold change (Supplementary Figure 1H). To verify RNA-seq data Ig expression, naïve, spleen-derived B cells from WT and KO mice were activated with LPS and IL4 and selected mRNAs were analyzed by northern blot. The IgGλ, IgGĸ transcripts are indeed less abundant and migrate faster on the gel in TENT5C KO compared to those from WT, confirming that they are TENT5C substrates (Figure 1H). Importantly, *TENT5C* is the only one of the four members of the *TENT5* gene family expressed at detectable levels in B cells and undergoing strong induction during their activation and differentiation as we measured by real-time quantitative PCR (qPCR) (Supplementary Figure 1I). Interestingly, its expression positively correlates with upregulation of PABPC1 what suggest that mRNA polyadenylation contributes to transcriptional reprogramming of the B cells response (Supplementary Figure 1I).

Finally, analysis of our ONT sequencing data revealed that transcripts with high translation rates significantly differ in their 3’ end polyadenylation status. Immunoglobulin transcripts possess significantly longer poly(A) tails compared to other highly abundant mRNAs which is opposed to previous reports indicating that short poly(A) tails are a conserved feature of highly expressed genes (Lima et al., 2017). For example, in an activated B cells mRNAs encoding ribosomal proteins have significantly shorter poly(A) tails (with a median value 50 bp) than immunoglobulin coding transcripts (with median values 80-120 bp) which are also highly expressed genes (Supplementary Figure 1H).

### B cells isolated from TENT5C KO produce less antibodies

Next, we analyzed the impact of inefficient transcript polyadenylation on immunoglobulin production at the protein level using western blot. TENT5C KO B cells grown *in vitro* produced fewer antibodies than those isolated from WT littermates (both light and heavy chains) while levels of other secreted proteins: interleukin 6 (IL-6), as well as ER-associated chaperonin – GPR94 were not changed as much (Figure 2A). According to this, the analysis of media collected from B cell cultures confirmed a decreased level of secreted antibodies by activated TENT5C deficient B lymphocytes (Figure 2B). Moreover, flow cytometry revealed that the level of IgG1 positive-cells is significantly decreased in TENT5C deficient B cells after 3 days of activation with IL-4 and LPS *in vitro* (Figure 2C).

**Figure 2.**
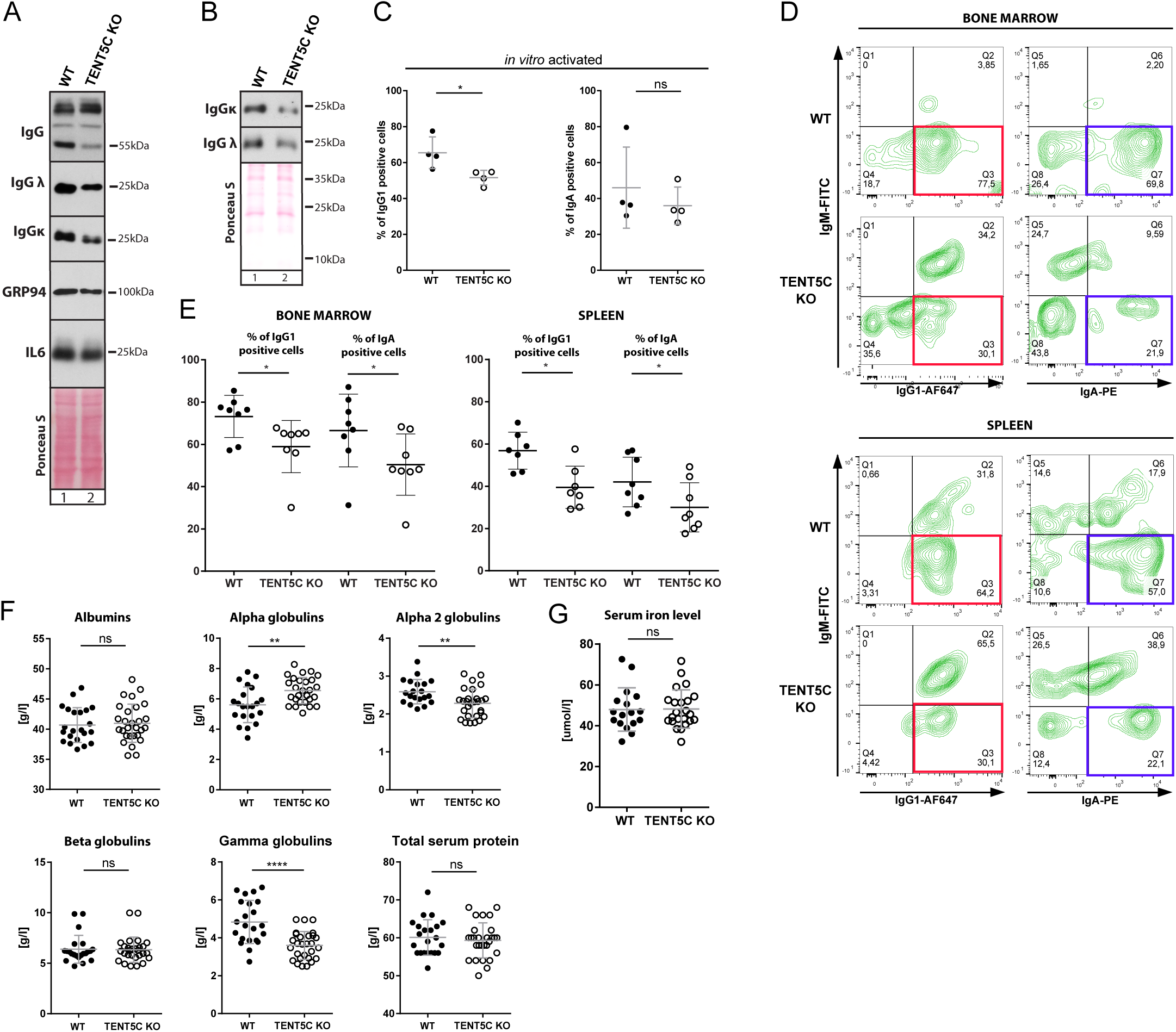
Polyadenylation of immunoglobulins mRNAs by TENT5C enhances their expression. (A) Western Blot analysis of IgG heavy and light (κ and λ) chains, IL6 and GRP94 from WT (lane 1) and TENT5C KO (lane 2) B cells, activated with LPS and IL4 for 3 days. Ponceau S staining was used as a loading control. (B) Western Blot analysis of immunoglobulins secreted to media by WT (lane 1) and TENT5C KO (lane 2), over 3 days of activation. Ponceau S staining was used as a loading control. (C) Percentage of IgG1 and IgA positive cells in WT and TENT5C KO B cells activated *in vitro* with LPS and IL4 for 3 days. P values were calculated using the Mann-Whitney U test (n =4). (D) Flow cytometry analysis of intracellular Ig isotypes levels in mature resting PCs isolated from bone marrow or spleen. Plasma cell populations were analyzed based on CD138, CD19, CD45R, IgM, IgG1 and IgA staining. See gating strategy in Supplementary Figure 4. IgG1-positive cells are marked with red rectangles, IgA-positive cells are marked with blue rectangles. (E) Percentage of mature resting PC positive cells for intracellular IgG1 or IgA isotypes. P values were calculated using the Mann-Whitney U test (n≥7). (F) Examination of blood serum albumins, alpha globulins, alpha 2 globulins, beta globulins, gamma globulins and total protein levels in TENT5C KO and control animals by SPEP. P values were calculated using unpaired t-test with Welch’s correction (n≥25). (G) Evaluation of iron level in blood serum of TENT5C KO and control animals. P value was calculated using unpaired t-test with Welch’s correction (n≥17).

Next, to evaluate the effect of these phenomena for the physiology of an organism, we assessed the intracellular (cytoplasmic) levels of IgG1 and IgA-positive cells in a population of CD138-positive cells isolated from bone marrow and spleen. The intracellular levels of immunoglobulins, in contrast to surface immunoglobulins, reflects secreting ability of these cells. As expected, the percentages of IgG1 and IgA-positive cells is lowered in TENT5C KO mice, which confirms altered immunoglobulin production (Figure 2D, E). To extend these studies, we applied blood serum electrophoresis to compare globulin fractions from WT and KO mice. In agreement with previous *in vitro* and *in vivo* cell analyses, we have found a substantial decrease of gamma globulin fraction (which contains mainly whole antibodies) in KO mice plasma reflecting alterations in their production and secretion by B cell lineage (Figure 2F). The effect was specific since total serum protein concentrations, as well as albumin, alpha and beta globulin plasma sub-fractions, were not changed. A slight decrease of alpha 2 globulin concentration in TENT5C KO serum is most likely a consequence of previously reported by us microcytic anemia (Mroczek et al., 2017) that develops in KO mice presumably as a result of inhibited globin synthesis, but not iron uptake deficiency as its levels were unchanged in the serum (Figure 2G).

In sum, we conclude that TENT5C KO leads to decreased immunoglobulin synthesis in mice.

### TENT5C is mainly expressed at late steps of B cell differentiation

Next, to take advantage of cytometric techniques in further analyses of B cell lineages, we have generated a TENT5C-GFP *knock-in* mouse. Similarly to the previously published TENT5C-FLAG (Mroczek et al., 2017), the *knock-in* GFP mouse line did not display any gross phenotype. Subsequently, flow cytometry analyses of *in vitro* activated B cells isolated from TENT5C-GFP mice revealed a distinct GFP-positive cell population, absent in the non-tagged controls confirming the utility of our mouse model (Figure 3A). Naïve B cells from TENT5C-GFP mice and WT littermates were activated with LPS and IL-4 and subjected to cytometric analysis to measure TENT5C-GFP and CD138 plasma cell markers levels in a time-dependent manner. This analysis revealed that TENT5C is mainly expressed in the population of CD138-positive cells what suggests the involvement of this enzyme at the last steps of B cell differentiation (Figure 3B). To confirm this hypothesis, we have systematically examined spleen and bone marrow-residing B cell and plasma cell subpopulations from young adult (12-15 weeks) unimmunized WT and TENT5C-GFP mice using multicolor flow cytometry. The GFP-tag *knock-in* does not affect the general distribution of B cell and plasma cell populations. However, this approach revealed that the CD138^high^ B cell subset is also highly GFP-positive in both BM (up to 76%) and spleen (up to 93%) (Figure 3C). In turn, GFP-positive cells were not detected in B cell subsets at early stages of differentiation (Supplementary Figure 2). The detailed gating strategy is presented in Supplementary Figure 3 and 4. Thus TENT5C-GFP is mainly expressed in the last stages of B cell differentiation as revealed by detailed plasma cell subpopulation analyses including dividing plasmablast (CD19^high^/CD45R^high^), early plasma cells (CD19^high^/CD45R^low^) and mature resting plasma cells (CD19^low^/CD45R^low^) (Figure 3D and Supplementary Figure 4). Finally, we confirmed those results with immunostaining of spleen sections showing that GFP-fused TENT5C is mainly expressed in CD138-positive cells (Figure 3E).

**Figure 3.**
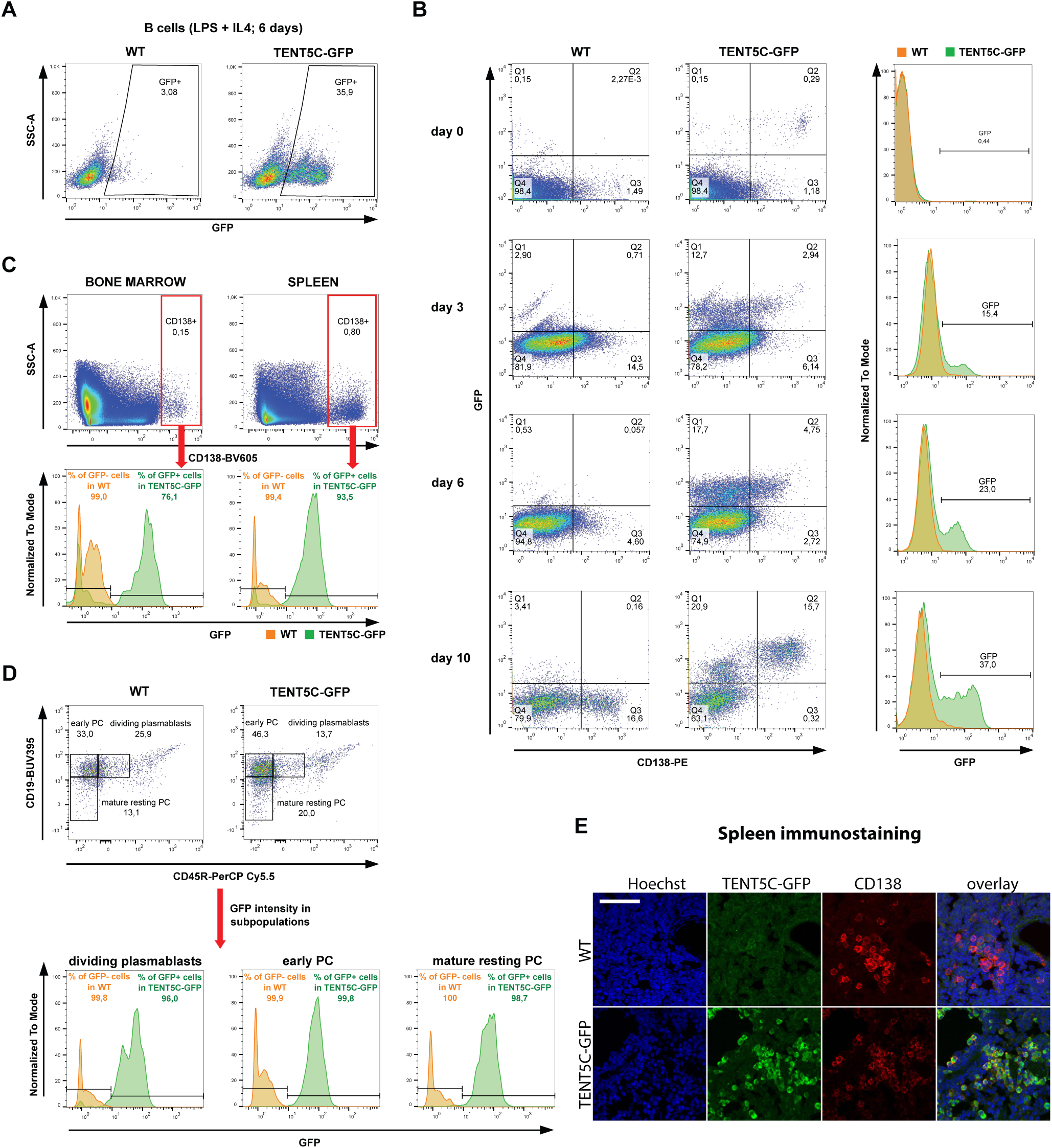
Expression of TENT5C is limited to the last stages of B cell lineage differentiation. (A, B) Flow cytometry analysis of TENT5C-GFP expression in a subpopulation of CD138^high^ cells activated *in vitro* with LPS and IL4 for 6 days (A) or up to 10 days (B) presented as pseudo-color dot blot (A) and/or histograms of GFP-fluorescence intensity (B). Grey color refers to WT, green is related to TENT5C-GFP. Detailed gating strategy is presented in related Supplementary Figure 4. (C, D) Flow cytometry analysis of TENT5C-GFP expression in splenocytes and BM (C) and different splenic PC subpopulations (D): dividing plasmablasts, early PC, mature resting PC. Colour code as described above. See also Supplementary Figures 2 and 4. (E) Immunohistochemical staining for plasma cell marker CD138 and GFP in spleens of wild-type and TENT5C-GFP *knock-in* mice. Scale bar denotes 50 μm.

### TENT5C expression is stimulated by innate signaling

Next, to define whether other stimuli than a combination of LPS and IL-4 lead to TENT5C upregulation, we have tested main types of B cell activation. Naïve B cells isolated from TENT5C-GFP mice and WT littermates as a control were activated with a panel of agonists of TLR receptors, mainly pathogen-associated molecular pattern (PAMP) molecules, ligands of CD40 (T-cell-dependent signaling) and BCR (B cell receptor). Subsequent flow cytometry analyses of those cells performed in a time course experiment revealed a significant number of GFP and CD138-positive cells, similar to positive control, as a result of stimulation of selected TLR receptors, including TLR1/2 (Pam3CSK4), TLR2 (HKLM), TLR4 (LPS, *E. coli* K12), TLR6/2 (FSL1) and TLR9 (ODN1826) (Figure 4). Stimulation of TLR3 (low and high molecular weight Poly(I:C)), TLR5 (Flagellin *S.typhimurium*) and TLR8 (ssRNA40/LyoVec) showed a rather limited effect on TENT5C-GFP expression.

**Figure 4.**
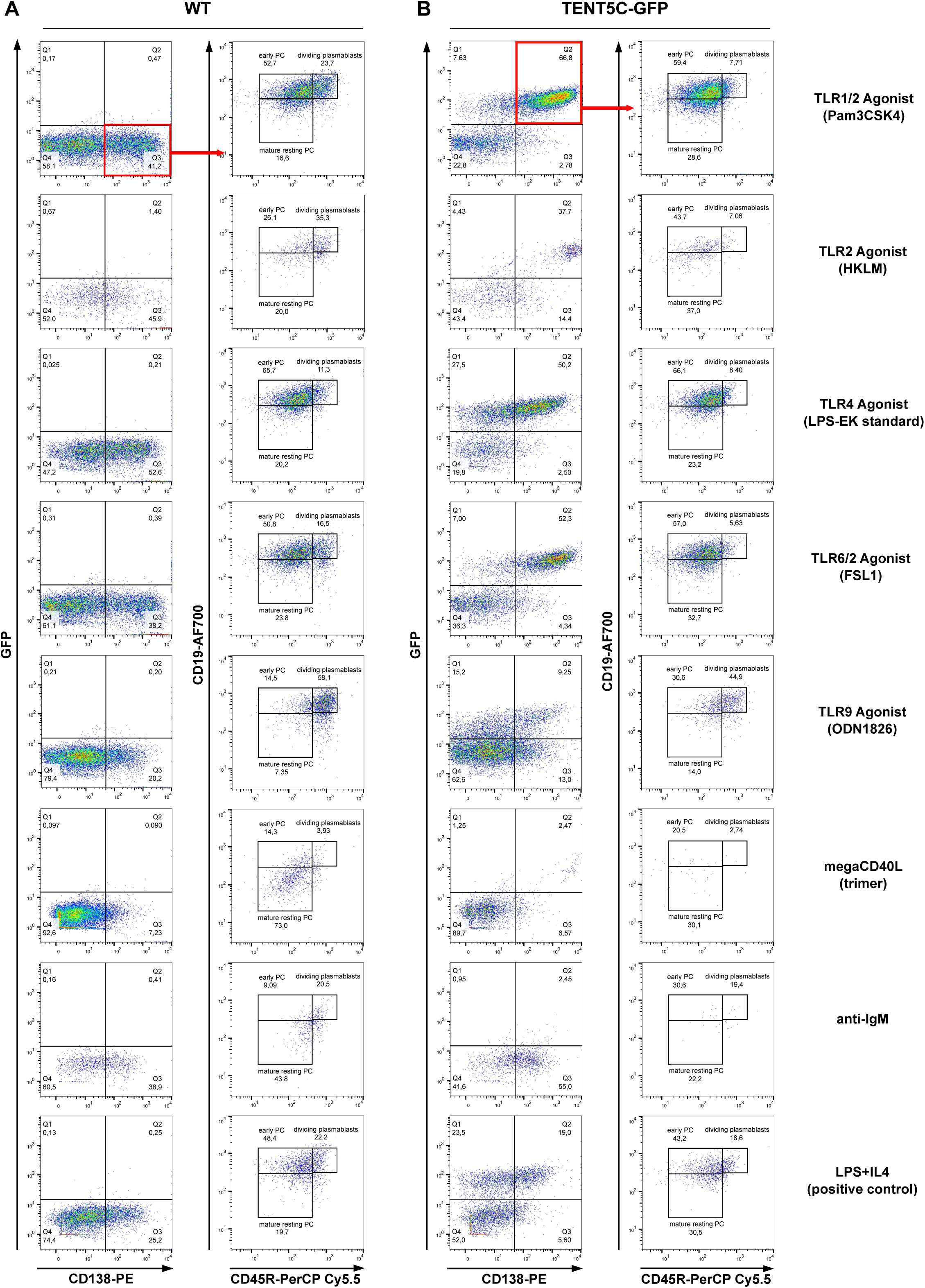
TENT5C is specifically upregulated by specific TLR signaling. (A, B) Analysis of the B cells activation process, measured by the evaluation of co-expression of CD138 and GFP molecules (red quarter, left columns) and further differentiation process of CD138^high^GFP^pos^ into dividing plasmablasts, early PC and mature resting PC (right column). B cells isolated from WT (A) and TENT5C-GFP (B) mice, were treated with different activators (indicated on the right part of the plot) for 4 days: TLR1/2 agonist (Pam3CSK4), TLR2 Agonist (HKLM), TLR4 agonist (LPS-EK standard), TLR6/2 agonist (FSL1), TLR9 Agonist (ODN1826), megaCD40L (trimer), anti-IgM and LPS/IL4 as a positive control. Flow cytometry analysis was based on the live/dead, CD138, CD45R and CD19 staining and additionally GFP. See also Supplementary Figure 4.

Subsequent analysis of the plasma cell subpopulations in CD138^high^ (Q3 for WT; Figure 4A) and CD138^high^GFP^pos^ (Q2 for TENT5C-GFP; Figure 4B) cell fractions using CD19 and CD45R markers showed a similar response to the treatment for cells isolated from both mouse lines. In turn, signaling provided by BRC stimulated with polyclonal F(ab’)2 goat anti-mouse IgM and CD40 receptor with megaCD40L (trimeric variant) had limited effect on TENT5C expression, however, they induced the differentiation of plasma cells (Figure 4).

To assess whether antibody production is also impaired in TENT5C KO B cells activated via innate signaling pathways, we activated B cells from WT and KO with TLR agonists with the highest effect on TENT5C expression (TLR 1/2, TLR4, TLR6/2 and TLR9 agonists) in a time course, then collected cells and medium for western blot analysis at day 3 and 5. Indeed, we observed decreased secretion of both heavy IgG and light chains in KO, while α-tubulin and GRP94 levels were unaffected (Supplementary Figure 5A, B).

All this together links B cell-intrinsic innate signaling via selected TLRs with the regulation of adaptive humoral immunity modulated by ncPAP TENT5C and reveals this enzyme as an important modulator of B cell response.

### TENT5C regulates B cell differentiation

TENT5C KO leads to an increased B cell proliferation rate, suggesting that TENT5C may control the process of their differentiation into plasma cells (Mroczek et al., 2017). Since the terminal differentiation of B cells takes place in secondary lymphoid organs, we carried out extended phenotyping of B cells in the spleen as well as bone marrow. First, we observed that spleens in KO mice are about 20% enlarged compared to those isolated from WT (Figure 5A). The effect is specific since there is no difference in overall animal mass (Figure 5B). As this observation strongly suggested enhanced proliferation rates we have systematically examined B cell subpopulations from spleens and bone marrow of conventionally cohoused adult unimmunized littermates (12-16 weeks) using multicolor flow cytometry.

**Figure 5.**
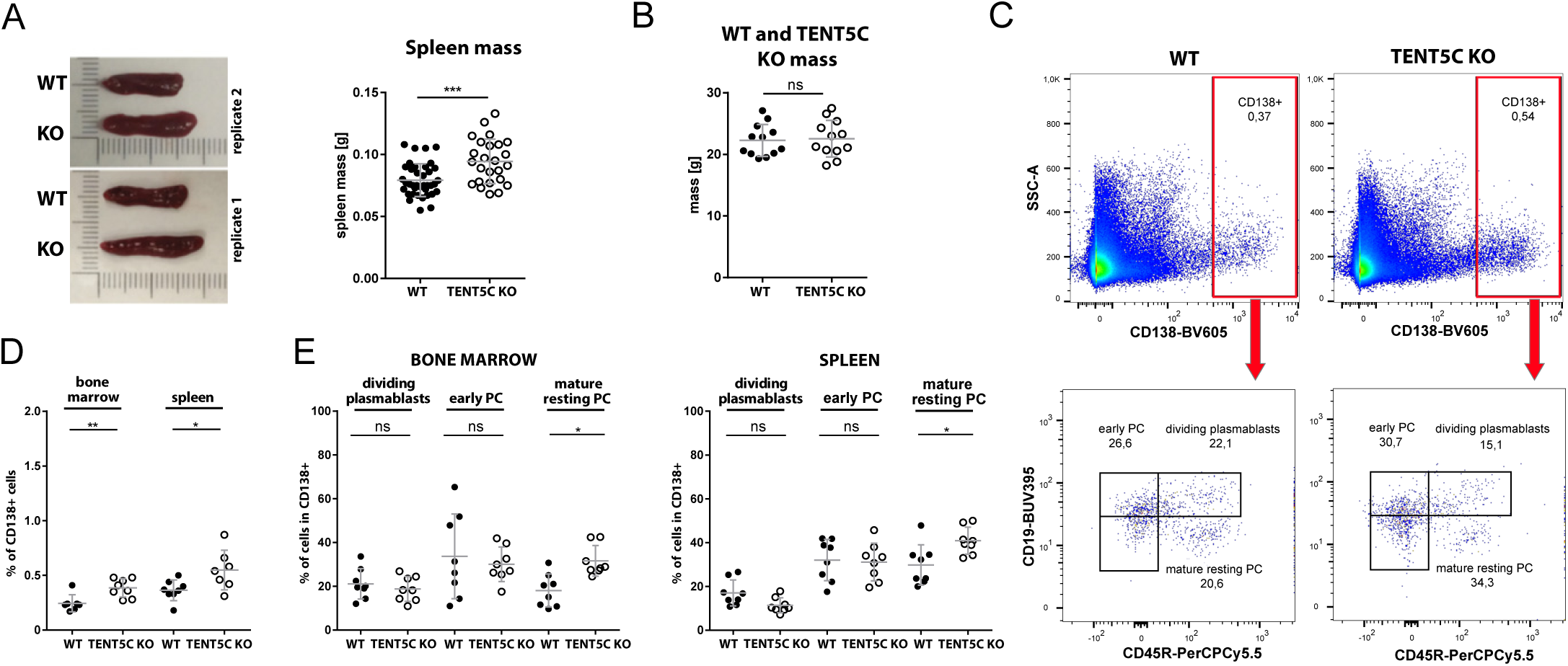
TENT5C negatively regulates B cell lineage differentiation into PC. (A) Comparison of spleen mass in WT and TENT5C KO mice (right panel). P values were calculated with unpaired t-test with Welch’s correction (n≥25). Pictures (left panel) show the representative difference in the spleen size between WT and TENT5C KO in two replicates. (B) Comparison of body mass in WT and TENT5C KO mice. P values were calculated with unpaired t-test with Welch’s correction (n=12). (C) Flow cytometry quantitative analysis of CD138^positive^ cells (upper pseudocolor dot blots) and particular CD138^high^ subsets: dividing plasmablasts, early PC and mature resting PC (lower pseudocolor dot blots) in WT and TENT5C KO. Plasma cell populations were analyzed based on CD138, CD19, CD45R, IgM, IgG1 and IgA staining. See gating strategy in Supplementary Figure 4. (D) Percentages of CD138^high^ cells in WT and TENT5C KO in both bone marrow and spleen. P values were calculated using the Mann-Whitney U test (n≥7). See Supplementary Figure 6 for other cell subpopulations. (E) Comparison of CD138^high^ subpopulations (selected as shown in the panel C): dividing plasmablasts, early PC, mature resting PC in WT and TENT5C KO in both bone marrow and spleen. P values were calculated using two-way ANOVA with post-hoc Bonferroni test (n=8).

Interestingly, we observed that the number of CD138^high^ cells in spleen and bone marrow was significantly increased in TENT5C KO mice (Figure 5C-D). Next, we carried out the quantitative determination of CD138^high^ plasma cell subpopulations in WT and KO mice. This has revealed mature resting PCs as the only ones whose number is increased in TENT5C KO mice spleen and bone marrow while numbers of dividing plasmablasts and early PCs were slightly decreased or not changed in KO (Figure 5E). Interestingly, other B cell subpopulations in bone marrow (pre-proB, pro-B, pre-B, immature, early/late mature, transitional B) and splenic (transitional (T1/T2/T3), marginal zone (MZP & MZ), follicular (I & II)) were not affected by TENT5C KO (Supplementary Figure 6). All these observations clearly suggest that the lack of TENT5C enhances B cell proliferation and differentiation *in vivo* and confirms previous *in vitro* findings.

Next, we asked whether TENT5C shapes the secondary antibody repertoire generated by class-switch recombination (CSR), which replaces IgM with other isotypes during B cell differentiation into plasma cells, and whether it differs in the KO compared to the WT. The changes in the class profile of presented antibodies may indicate a disturbance in the CSR process. We observed that CD138^high^ B cell subsets in TENT5C KO lose IgM expression much faster as compared with WT which confirms their faster proliferation and accelerated selection of IgG1-expressing polyclonal plasma cells (Figure 6A, B, D, E). Our analysis also shows that in comparison to WT mice there are less IgA-positive plasmocytes in TENT5C KO mice, indicating problems with class switch recombination (CSR) in the KO mutant (Fig 6B, C, F). The general concentration of immunoglobulins is lowered in KO cells as a result of a diminished expression level and they accumulate membrane-bound IgG1 (Figure 6E). This is in agreement with the results we obtained for *in vitro* cultured B cells (Figure 2A, B).

**Figure 6.**
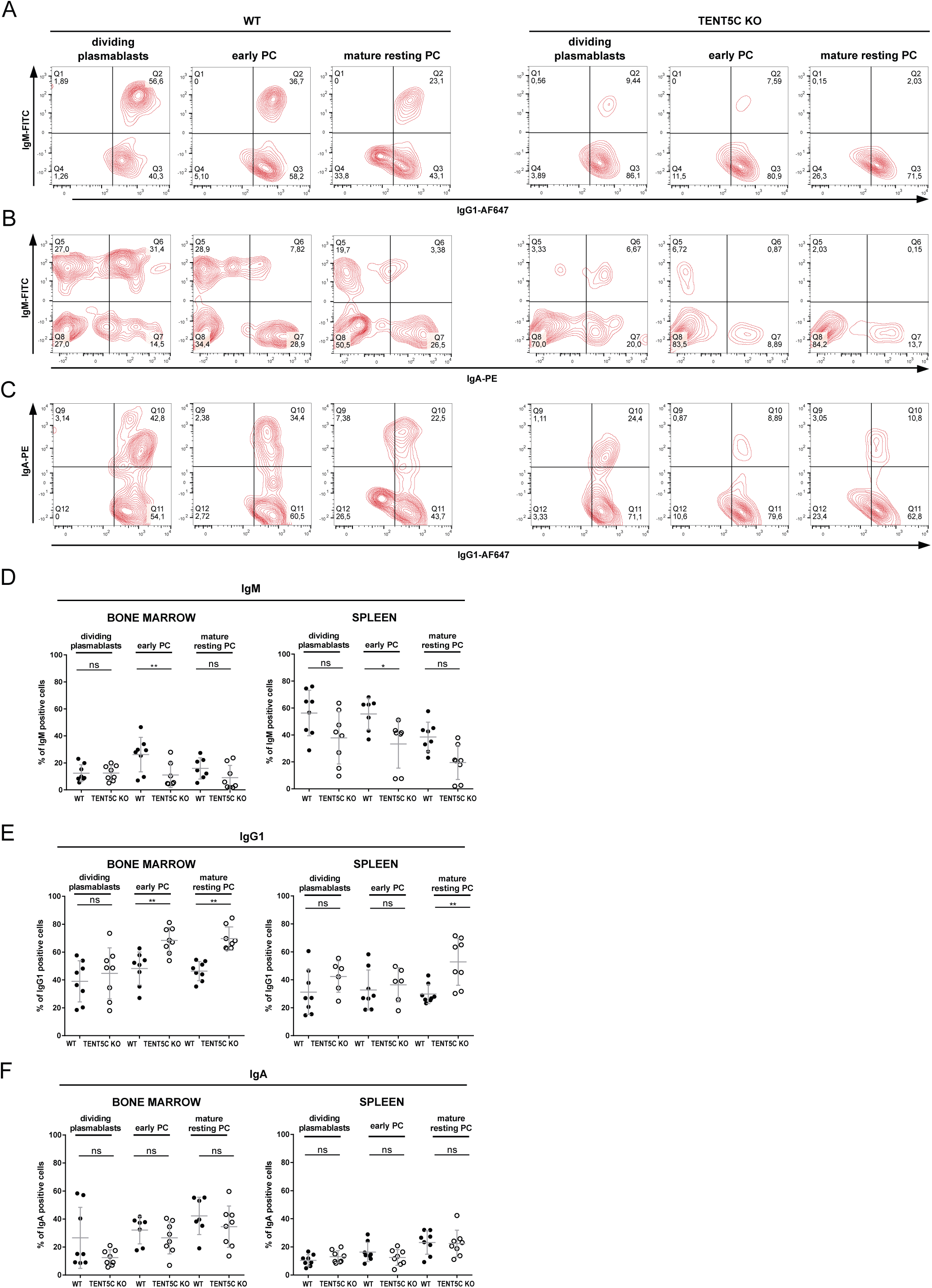
TENT5C KO plasmacytes are characterized by the abnormalities in the class switch recombination (CSR). (A) (B) (C) Flow cytometry analysis of surface Ig isotypes: IgM vs IgG1 (A), IgM vs IgA (B), IgA vs IgG1 (C) in dividing plasmablasts, early PC and mature resting PCs isolated from bone marrow or spleen (representative contour blots are shown). Plasma cell populations were analyzed based on CD138, CD19, CD45R, IgM, IgG1 and IgA staining. See gating strategy in Supplementary Figure 4. (D, E, F) Percentage of cells positive for surface IgM (D), IgG1 (E), or IgA (F) immunoglobulins in dividing plasmablasts, early PC and mature resting PCs isolated from bone marrow or spleen. P values were calculated with two-way ANOVA with post hoc Bonferroni test (n=8).

### TENT5C is an ER-associated protein shaping for ER functionality

The transition of B cells into immunoglobulin-secreting plasma cells requires a significant expansion of secretory organelles, given that ER-specific chaperones and folding enzymes facilitate the post-translational structural maturation of Igs. Since TENT5C modifies transcripts encoding Igs in responding B cells it may have a possible ER-related function. To examine the link between TENT5C activity and ER expansion during B cell responses, we performed a fractionation of activated B cells isolated from TENT5C-FLAG mice followed by western blot analysis. This revealed a significant fraction of the membrane-bound enzyme, which strongly suggests TENT5C ER-association (Figure 7A). This result was confirmed by a partial intracellular co-localization of endogenous TENT5C-GFP protein with the selectively stained endoplasmic reticulum (ER) in isolated CD138^pos^ cells (Figure 7B). Finally, an ER-related function is supported by co-immunoprecipitation (Co-IP) experiments using high-affinity anti-GFP nanobodies followed by high-resolution mass spectrometry (MS) which revealed that TENT5C-GFP interacts with ribosomal proteins; thus it may directly polyadenylate immunoglobulin mRNAs at the rough ER (RER) (Supplementary Dataset 3).

**Figure 7.**
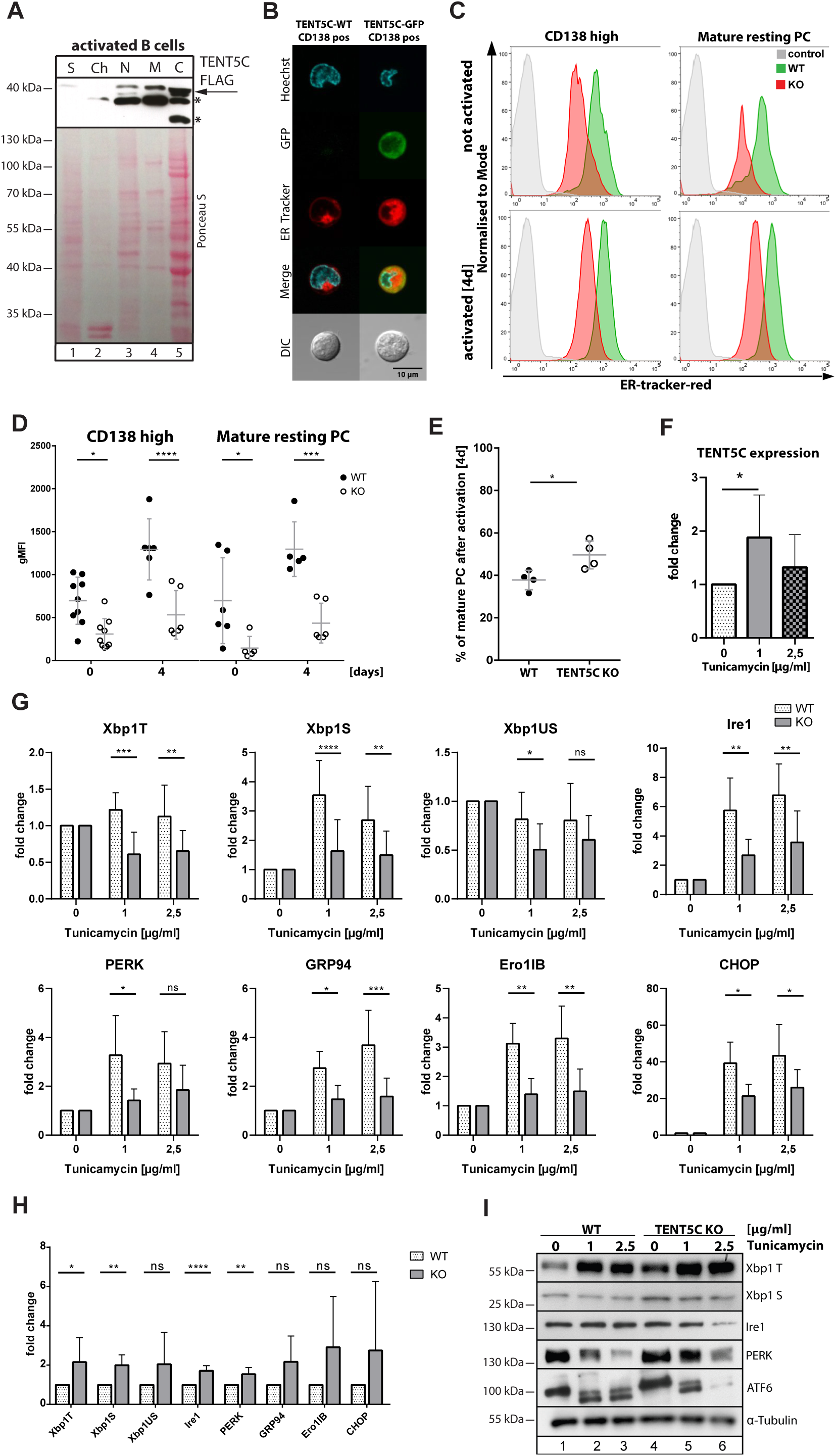
TENT5C is a membrane-associated protein and affects UPR response. (A) Endogenous TENT5C localizes mainly to the membranes and cytosolic fractions. B cells were isolated from TENT5C-FLAG mouse, activated with LPS and IL4 for 3 days and fractionated for S – cytoskeletal (lane 1), Ch – chromatin (lane 2), N – soluble nuclear (lane 3), M – membrane (lane 4), C – cytoplasm (lane 5), and analysed with Western blot using an anti-FLAG antibody. Arrow indicates the position of TENT5C-FLAG while asterisks indicate nonspecific bands detected by anti-FLAG antibodies. (B) Staining of the nucleus and the ER of CD138 positive cells isolated from wild-type and TENT5C-GFP *knock-in* mice. Scale bar denotes 10 µm. (C) Flow cytometry analysis of ER-fluorescence intensity in CD138^high^ cells and mature resting PC. Cells were isolated from WT (marked in green) and TENT5C KO (shown in red). The measurements were performed at day 0 and 4 days after activation (LPS and IL4). Plasma cell populations were analyzed based on CD138, CD19, CD45R, and ER-tracker. See gating strategy in Supplementary Figure 4. (D) Quantification of ER volume, measured as gMFI (geometric mean of fluorescence intensity) of ER-tracker, based on flow cytometry. gMFI for ER-tracker was checked for CD138^high^ cells (left panel) and mature resting PC (right panel). The measurement was done at day 0 and 4 days after activation (LPS and IL4). P values were calculated using two-way ANOVA with post-hoc Bonferroni test. (E) Percentage of mature PC 4 days after activation with LPS and IL4. P value was calculated with the Mann-Whitney U test (n=4). (F) TENT5C expression level after tunicamycin treatment. B cells from WT mice were isolated and activated with LPS and IL-4 for 3 days, then treated with tunicamycin in indicated concentration for 5 hours. The amount of TENT5C mRNA was checked by qPCR. P values were calculated using Student’s t-test (n≥6). (G) qPCR analysis of UPR markers expression in WT and TENT5C KO cells after ER stress induction with tunicamycin. B cells from WT and TENT5C KO mice were isolated and activated with LPS and IL-4 for 3 days, then treated with tunicamycin in indicated concentration for 5 hours. Bars represent mean fold change values ±SD (n=6), P values were calculated using two-way ANOVA with post-hoc Bonferroni test. (H) qPCR analysis of basal ER stress level in B cells WT and TENT5C KO in 3^rd^ day after activation with LPS and IL-4. P values were calculated using Student’s t-test. (I) Western blot analysis of UPR markers (Xbp1 T – total, S – spliced, Ire1, PERK and ATF6) in WT and TENT5C KO cells after ER stress induction with tunicamycin. The α-tubulin was used as a loading control.

As B lymphocyte maturation requires a significant increase in ER volume and TENT5C affects the rate of the differentiation process we compared its size and expansion dynamics during the activation of WT and KO B cells using specific ER-tracker dye labeling followed by flow cytometry analyses. The results of these experiments showed that a lack of TENT5C impairs the capacity of the secretory pathway through the reduction of ER volume (naïve and mature resting PC) (Figure 7C, D). Moreover, the dynamic of the ER expansion after activation is much slower in the TENT5C-deficient cells despite their accelerated differentiation into plasmocytes (Figure 7E). Reduced ER is consistent with a decreased level of both immunoglobulin encoding transcripts and the main chaperone Hspa5 (BIP), necessary for the correct functioning of the ER in TENT5C KO cells (Supplementary Dataset 2). Thus, our findings together with the fact that the overall level of antibody production in KO cells is reduced, strongly suggests that B cells isolated from TENT5C KO may have reduced ER stress levels. In order to analyze the ER-stress response, we treated activated WT and KO B cells with the standard ER stress-inducing agent tunicamycin (Tu) and then analyzed unfolded protein response (UPR) markers with qPCR and western blots. Interestingly, induction of ER-stress enhances TENT5C expression (Figure 7F). Surprisingly, the initial ER stress level is enhanced in the mutant compared to WT however the KO cell response for Tu treatment was significantly diminished as shown by Xbp1 mRNA splicing and expression of selected markers IRE1, PERK, GRP94, CHOP, Ero1-LB, showing a general downregulation of unfolded protein response (Figure 7G-I).

Concluding, TENT5C dysfunction leads to a reduced ER volume and capacity of the ER stress response as a probable consequence of a decreased load of Ig.

## Discussion

B cell development in mice and humans has been extensively studied, revealing complex physiological changes, driven by different signaling pathways, which influence the genome (somatic hypermutation, class-switch recombination), transcriptome (through the coordinated action of transcription factors) and proteome (ER reorganization, post-transcriptional gene expression regulation) in differentiating cells. In this study, we provide evidence for cytoplasmic polyadenylation, driven by TENT5C, being the previously undescribed mechanism involved in the regulation of immunoglobulin expression and B cell differentiation. Our data indicate that the role of cytoplasmic polyadenylation is broader than previously anticipated and provides a new layer to the regulation of immunoglobulin expression.

Recently, we presented the first experimental data for a new family of non-canonical cytoplasmic poly(A) polymerases TENT5 (formerly FAM46) (Warkocki et al., 2018). One of the members, TENT5C, was shown to be a specific growth suppressor in multiple myeloma cells (Mroczek et al., 2017; Zhu et al., 2017) and is the only TENT5 family member expressed at significant levels in B lymphocytes. In this work, using direct RNA sequencing by Oxford Nanopore Technologies, we identified immunoglobulin mRNAs as TENT5C specific targets in activated B cells. To our knowledge, this is also the first report showing a global view of poly(A) tails in B cells. Our approach offered high-quality, full-length mRNA sequences, including UTRs and poly(A) tails giving deeper insights into transcriptome shaping/regulation comparing to classical RNA-seq experiments. Comparing to the currently used RNA 3’-end research techniques such as TAIL-seq, PAL-seq, TED-seq, PAC-Seq or recently FLAM-seq (Chang et al., 2014; Harrison et al., 2015; Legnini I., 2018; Nicholson and Pasquinelli, 2018; Welch et al., 2015; Woo et al., 2018) it is characterized by a relative technical simplicity, and in contrast to other RNA-seq methods, no PCR-biases are introduced into libraries.

Despite the recent dynamic expansion of RNA 3’-terminome research, little was known how cytoplasmic ncPAP enzymes contribute to gene expression programs since such techniques were never applied for KOs of individual enzymes in physiological conditions. Cytoplasmic adenylation was mostly studied in the context of gametogenesis or in other instances in which transcription is arrested or spatially and temporarily separated. Importantly, even a ∼20% decrease of poly(A) tail length in TENT5C KO mice significantly diminishes a steady state concentration of immunoglobulins in the plasma or secreted by activated B cells cultured *in vitro.* This very strongly suggests that cytoplasmic polyadenylation is not restricted to deadenylated maternal mRNAs as in the case of gametogenesis. Since TENT5C is enriched at the ER and Igs are the main secreted proteins in B cells, we suggest that this may partially explain its specificity for immunoglobulins (Mroczek et al., 2017). However, as we were unable to identify any highly enriched sequence motif in TENT5C substrates, a more detailed analysis is needed to decipher the mechanism of its substrate specificity. Surprisingly, completely opposite to previous reports suggesting that highly expressed transcripts possess rather short poly(A) tails (Lima et al., 2017), we have shown that immunoglobulin transcripts, constituting the vast majority (up to 70%) of the PC transcriptome, have rather long poly(A) tails compared to other highly translated transcripts. It was previously shown that immunoglobulin regulation in a differentiation-dependent manner of membrane-associated or secreted IgM isoforms occurs by alternative polyadenylation at 3′ pre-mRNA, which is initiated by the cleavage stimulation factor CstF-64. (Enders et al., 2014; Peng et al., 2017; Pioli et al., 2014; Takagaki and Manley, 1998). Additionally, the regulation of switching from mIgH to sIgH in plasma cells is mediated by PABPC1 recruiting hnRNPLL to 3’-end of IgG transcripts in plasma cells making poly(A) tail a key mRNA feature for immunoglobulin production (Peng et al., 2017). Our finding confirms that adenylation is a key factor regulating immunoglobulin expression and for the first time provides evidence for cytoplasmic adenylation of immunoglobulin transcripts.

TENT5C was reported as one of the genetic signatures in ASC and a potential regulator of B cell differentiation and is one of the top 50 upregulated genes in spleen and bone marrow plasma cells (Shi et al., 2015). Our studies confirm a strong correlation of TENT5C expression with B cell proliferation and differentiation into plasma cells. Moreover, it is positively correlated with the upregulation of PABPC1 previously identified as TENT5C interactor in MM cells (Mroczek et al., 2017). Signaling from both surface (TLR1, 2, 4, 6) and intracellular (TLR9) TLR receptors strongly upregulate TENT5C levels thus promote the B cell lineage differentiation and enhance immune response, however detailed dissection of TLR downstream signaling pathways require further investigation (Pasare and Medzhitov, 2005). Interestingly mice devoid of MyD88 (myeloid differentiation primary response gene 88) gene, which is one of the key elements of TLR signalling, reveal similar phenotypes to TENT5C deletion including: decreased steady-state levels of total serum immunoglobulins, decreased antigen-specific IgM and IgG1 antibody responses and abolished IgG2 antibody response in immunized mice (Kang et al., 2011). This strongly suggests that TENT5C is one of the TLR signaling effectors in B cells. In agreement, TENT5C upregulation by specific B cell innate signaling is underlined by the fact that it is not affected by the stimulation of BRC and CD40 receptor (T cell-dependent).

The transition of naïve B cells into ASC requires significant ER membrane expansion, given that the structural maturation of immunoglobulins is facilitated by ER-residing chaperones and folding machinery (Braakman and Hebert, 2013; Kirk et al., 2010; Liu and Li, 2008; Wiest et al., 1990). The increase in the volume of secretory organelles occurs through the generation of ER sheets and requires UPR signaling (Schuck et al., 2009). B cells activate all three branches of the UPR response including specific stress sensors: inositol-requiring enzyme-1α (IRE1α), protein kinase R (PKR)-like endoplasmic reticulum kinase (PERK) and activating transcription factor 6α (ATF6α). It has been shown that Xbp1 and Blimp-1 transcription factors are required for plasma-cell development and they link B cell physiology with UPR response (Gass et al., 2004; Reimold et al., 2001; Tellier et al., 2016). Additionally, a functional analysis of ASC signature genes identified 30% of the transcriptome as related to UPR (Shi et al., 2015). Thus, downregulation of immunoglobulin expression by TENT5C has to impair the unfolded protein response (UPR), which plays a pivotal role in the differentiation of ASC. In agreement with this, basic ER stress in naïve B cells from TENT5C KO mice is enhanced which reflects their faster proliferation and differentiation into PC while ER expansion dynamics is reduced. This also shows that cytoplasmic adenylation by TENT5C, being the posttranscriptional regulator of immunoglobulin expression, may have a profound effect on different aspects of B cell physiology, pointing at its, previously missed, important role not only in cell homeostasis but even its organismal-level one.

In conclusion, this study identified ncPAP TENT5C as a new factor involved in the regulation of immunoglobulin production through ER-associated polyadenylation in responding B cell lineage in mice. Thus, we demonstrate the significance of cytoplasmic poly(A) tail homeostasis as an important regulatory level for B cell immune response.

## Acknowledgments

We would like to thank Dominika Nowis for critical reading of the manuscript, Aleksander Chlebowski for microscopy assistance, Dominik Cysewski for high resolution mass spectrometry analyses, Janina Durys for the editing of the manuscript, Marcin Szpila for assistance in coordination of animal house work and Andrzej Dziembowski laboratory members for stimulating discussions. The equipment from www.ibb.waw.pl/en/services/mass-spectrometry-lab was sponsored in part by the Centre for Preclinical Research and Technology (CePT), a project co-sponsored by European Regional Development Fund and Innovative Economy, The National Cohesion Strategy of Poland.

## Author Contributions

SM and AD developed and directed the studies, AB and SM carried out majority of the biochemical and cell line experiments, MKK performed all flow cytometry analyses, PSK performed all bioinformatics analyses, OG participated in localization studies and coordinated work of the animal house, BT performed tissue immunostaining, KK purified the EIF4E protein, JG prepared CRISPR reagents and genotyped mice, EB generated transgenic animals. SM and AD wrote the manuscript with a contribution of PSK, AB, OG and MKK.

## STAR★Methods

### Key Resources Table

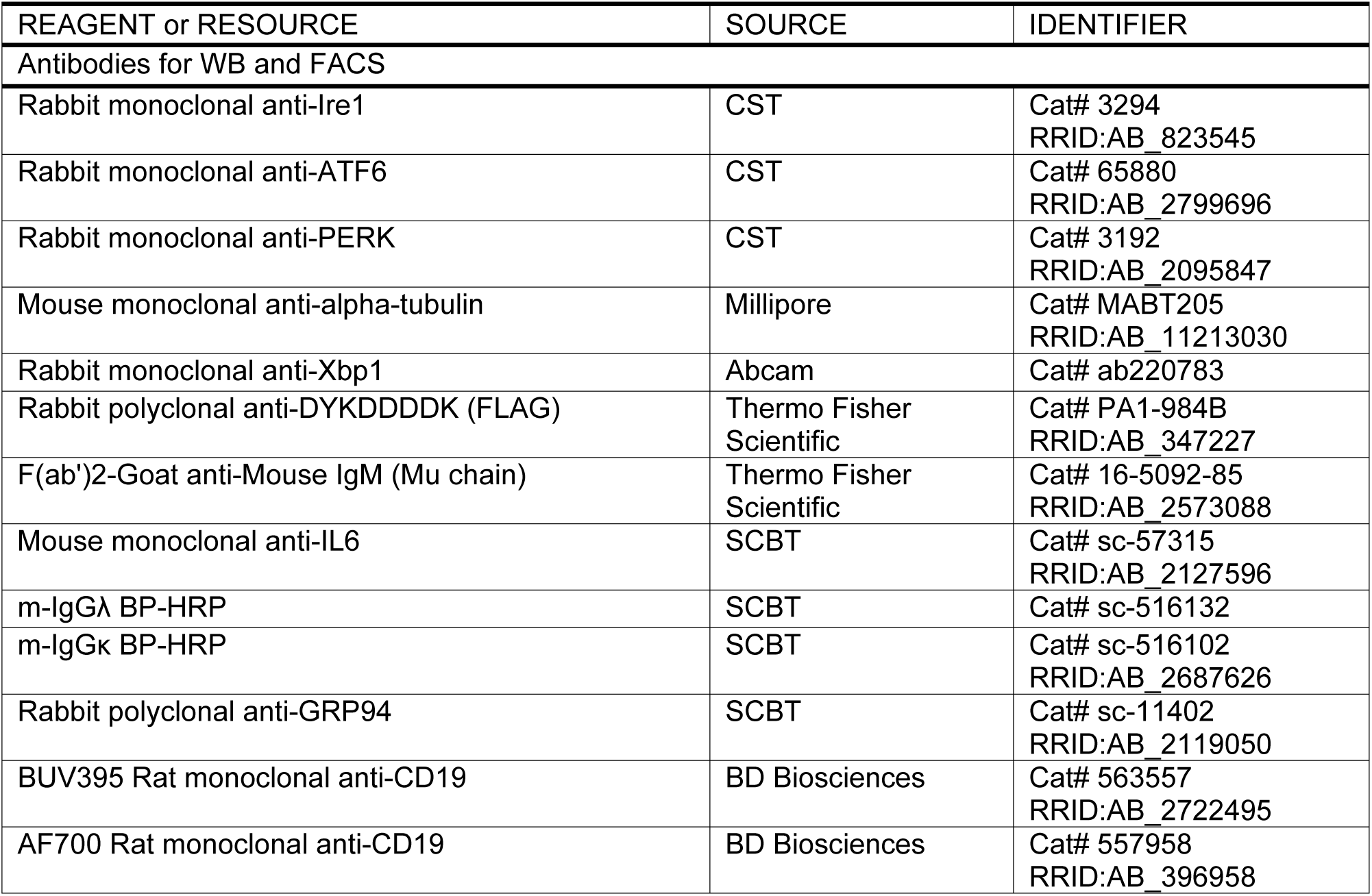
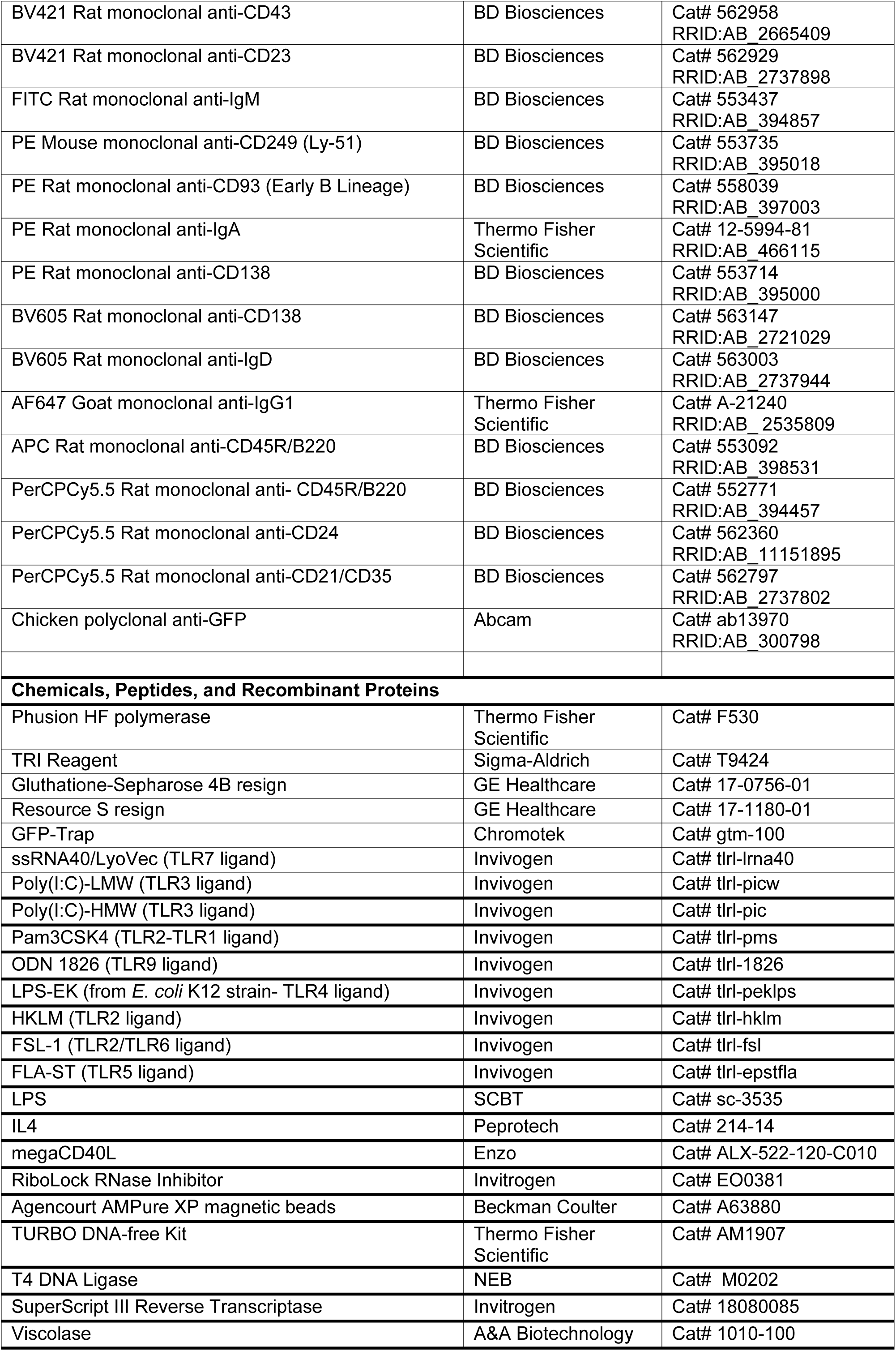
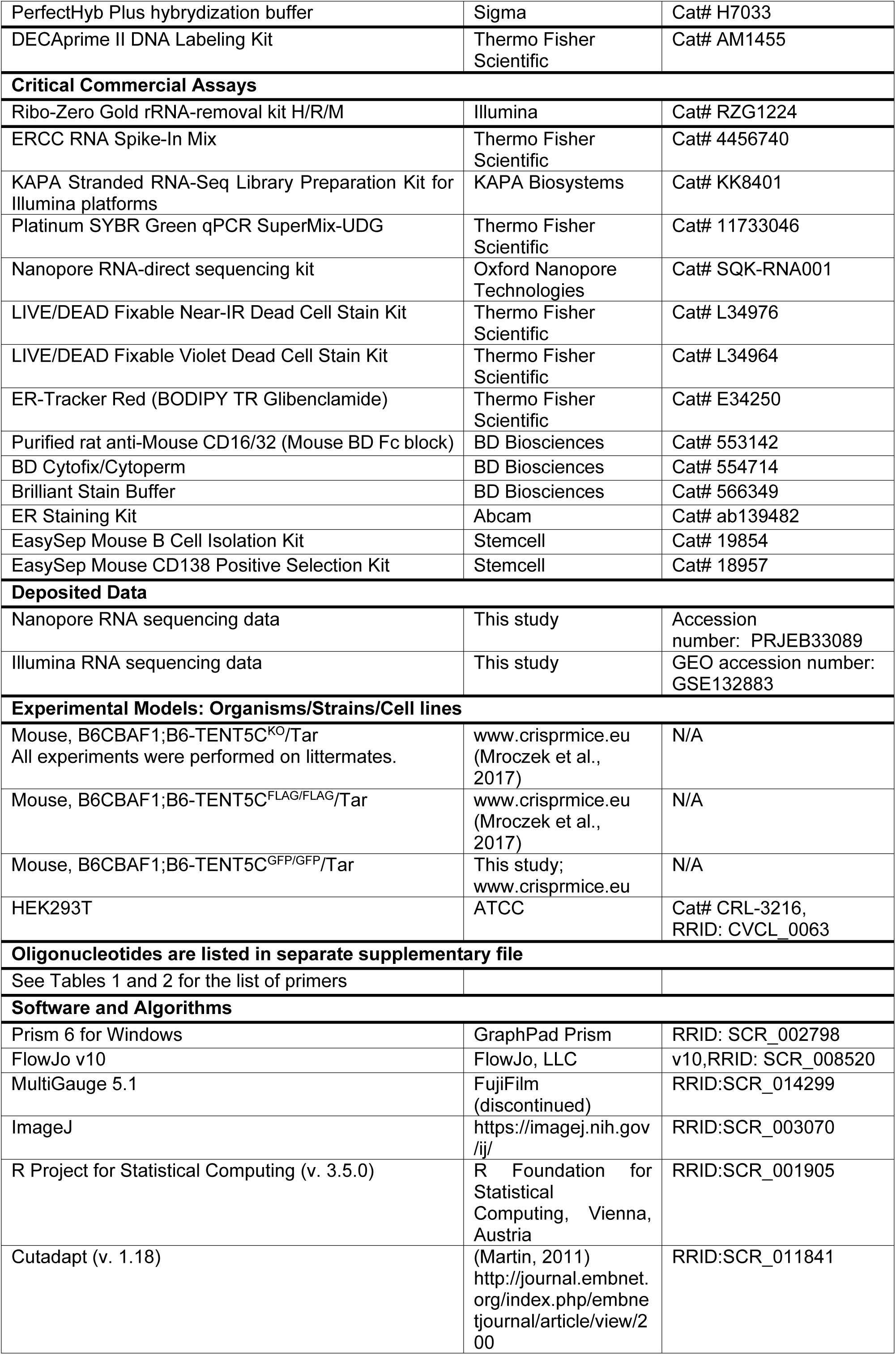
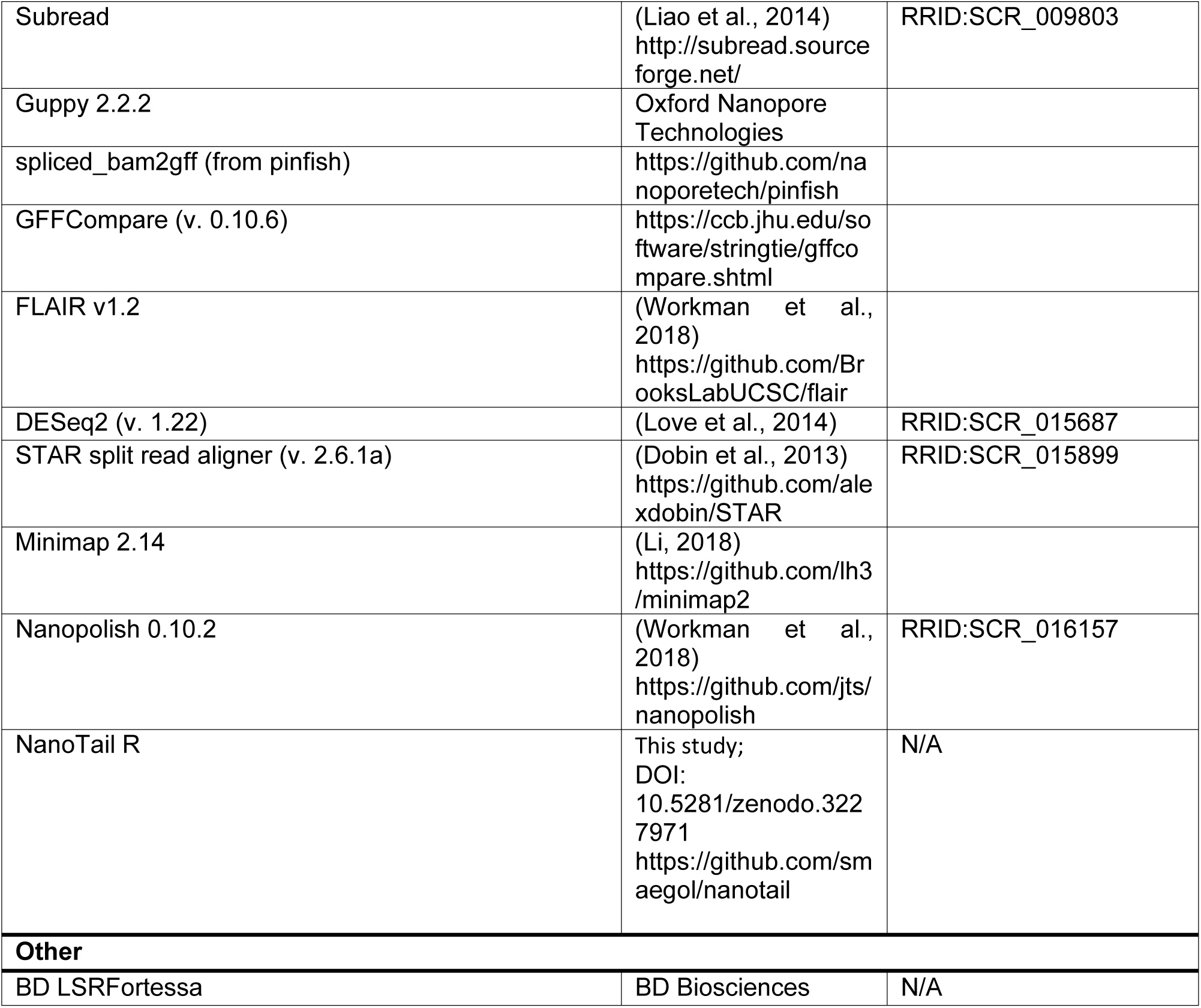

### LEAD CONTACT AND MATERIALS AVAILABILITY

Further information and requests for resources and reagents should be directed to and will be fulfilled by the Lead Contact, Andrzej Dziembowski.

### EXPERIMENTAL MODEL AND SUBJECT DETAILS

#### Mice

Mice were bred in conventional conditions at the Animal House at the Faculty of Biology, the University of Warsaw under 12h light/dark cycle at an ambient temperature of 22°C. Health monitoring was performed regularly at the IDEXX laboratory (reports are shown in Supplementary Table 3). Experimental mice originated from heterozygotic matings and as a result were cohoused littermates. All mice were sacrificed at age 12-16 weeks.

The mice used in this study were naive and had no previous history of experimentation or exposure to drugs. The TENT5C KO and TENT5C-FLAG mice strains were described previously (Mroczek et al., 2017). The TENT5C C-terminal-GFP *knock-in* animals were generated by the Mouse Genome Engineering Facility at the Institute of Biochemistry and Biophysics, Polish Academy of Sciences (https://crisprmice.eu/).

All procedures were approved by the I Local Ethical Committee in Warsaw.

### METHOD DETAILS

#### Generation of TENT5C C-terminal GFP Knock-in Mice

The TENT5C C-terminal-GFP *knock-in* mice line was generated using sgRNA (chimeric single-guide RNA) designed as close as possible to the STOP codon of TENT5C gene. The sgRNA was synthesized using T7 RNA polymerase and DNA template obtained with mTENT5C_GFP_sgRNA_F and Universal_gRNA_rev primers (underlined is T7 RNA polymerase promoter). All subsequent steps were performed as described previously (Mroczek et al., 2017). The dsDNA donor for in-frame C-terminal TEV-GFP *knock-in*, two 1kb homology arms flanking TENT5C STOP codon were amplified from C57BL/6J gDNA using mTent5C_TOPO-LF_1f/mTent5C_LF-TEV_1r and mTent5C_eGFP-RF_1f/mTent5C_RF-TOPO_1r primer pairs. TEV-eGFP coding sequence was amplified from pKK-TEV-eGFP (Szczesny et al., 2018) plasmid using TEV_1F and mCherry_GFP_1R primers.

Assembly of TEV-eGFP, 1kb homology arms and TOPO Zero Blunt (Thermo Fisher Scientific) cloning vector was performed with SLIC method and final pTOPO/F46C/TEV-eGFP construct was verified by sequencing. (Li and Elledge, 2012). Next, we amplified 865bp fragment of TENT5C-TEV-eGFP donor flanked with 60bp homology arms from pTOPO/F46C/TEV-eGFP and mTent5C-GFP_short_1F/ mTent5C-GFP-short_1R primers. PCR product was purified with AmpureXP beads (Beckman-Coulter) and stored at −20^°^C. Mice genotyping were carried out in PCR reaction using mTent5C_GFP_seqF and mTent5C_GFP_seqR primers and Phusion HotStart II Polymerase (Thermo) and gDNA isolated from of mice ears or tails fragments with HotShot method (Alasaad et al., 2008) or with Genomic Mini DNA isolation kit (A&A Biotechnology). The sequencing results were analyzed with Mutation Surveyor 4.0 (SoftGenetics). All primes used for mice generation are listed in Supplementary Table 1.

#### Tissue collection and blood analysis

Blood samples were collected terminally from the mandibular vein to EDTA or serum separator tubes. Complete blood count, gel electrophoresis of proteins and serum iron level analysis were performed at the Veterinary Diagnostic Laboratory LabWet in Warsaw (http://www.labwet.pl/) on the day of blood collection. For SPEP analyses were performed using SAS-MX SP-10 Kit (Helena-Biosciences). Serum samples were diluted in the buffer in a ratio of 1 to 4 and proteins were separated at a constant voltage of 80V through 25 min. Gels were quantified with Platinum software (Helena-Biosciences).

All mice were sacrificed by cervical dislocation. Spleen and femur and tibia bones were isolated immediately. Bone marrow was isolated using centrifugation method (Amend et al., 2016). BM was depleted of red blood cells using ACK lysis buffer (154.95 mM ammonium chloride, 10 mM potassium bicarbonate, 0.1 mM EDTA).

#### Primary cell culture and ex vivo B cell activation

Single cell suspension of splenocytes was obtained by mechanical tissue disintegration of the spleen through a 70 μm cell strainer. Then, splenocytes were additionally depleted from red blood cells using ACK lysis buffer before separation. Naïve B cells were isolated from spleen using immunomagnetic negative selection with EasySep™ Mouse B Cell Isolation Kit (Stemcell; 19854) and CD138^high^ cells were isolated from spleen and bone marrow with EasySep Mouse CD138 Positive Selection Kit (Stemcell; 18957) according to the manufacturer’s instructions.

Primary cells were cultured in RPMI 1640 ATCC’s modified (Invitrogen) supplemented with 15% FBS (Invitrogen), 100 nM 2-mercaptoethanol (Sigma), penicillin/streptomycin (Sigma) and activators or mitogens depending on the experiment: 20 ng/ml IL-4 (Peprotech), 0.5 μg/ml megaCD40L (Enzo), 10 μg/ml anti-IgM (Invitrogen), 20 μg/ml LPS (Santa Cruz), 1 μg/ml Pam3CSK4 (Invivogen), 10^8^ cells/ml HKLM (Invivogen), 10 μg/ml Poly(I:C), HMW and LMW (Invivogen), 10 μg/ml LPS-EK standard (Invivogen), 1 μg/ml FLA-ST (Invivogen), 100 ng/ml FSL1 (Invivogen), 1 μg/ml ssRNA40/LyoVec (Invivogen), 1 μM ODN1826 (Invivogen).

#### Spleen histology and plasma cell microscopy

For the histology of the spleen, animals were overdosed with ketamine/xylasine and perfused transcardially at 10 ml/min flow rate with PBS for 1min following 4% PFA in phosphate buffer (PB) for 2min at room temperature. Organs were dissected, post-fixed in PFA for 2hr at room temperature and suffused with 30% sucrose solution in PB overnight at 4°C. For immunohistochemically staining 10 μm-thick sections were cut with the cryostat, endogenous peroxidase was quenched with 3% H2O2 in TBS and sections were blocked 2 hrs with 10% rabbit serum and 1% BSA in TBS with 0.3% Triton X-100. Sections were incubated with primary antibodies anti-GFP from chicken (ab13970, Abcam, Cambridge, UK) and anti-CD138-PE from rat (BD Pharmingen, San Jose, CA, US) diluted 1000x and 500x, respectively, in blocking solution and developed with goat anti-chicken-Alexa-488 with Hoechst diluted in blocking solution. After quenching and incubation with antibodies sections were washed with TBS with Triton-X100 0.025% 3x 5min.

Isolated CD138^high^ cells were stained using ER Staining Kit (ab139482) according to the manufacturer’s instruction with following exceptions: Red Detection Reagent was diluted 2000x, Hoechst 33342 Nuclear Stain was diluted 500x and the staining time was shortened to 4 minutes.

The plasma cells and spleen section imaging was performed using a confocal system (Fluoview FV1000) equipped with a spectral detector (Olympus) and with 60x oil objective with 1.40 aperture. Images were processed using ImageJ software.

#### GST-eIF4E^K119A^ purification

Chemocompetent *Escherichia coli* BL21-CodonPlus-RIL strain (Stratagene) was transformed with plasmid pGEX-4T-3 carrying GST-eIF4EK119A. Cells were pre-incubated in standard Luria-Broth (LB) medium (with 0.1 mg/ml ampicillin and 34 mg/ml chloramphenicol) overnight and then transferred to Auto Induction Media Super Broth Base Including Trace Elements (Formedium) supplemented with 2% glycerol, kanamycin (50 mg/ml) and chloramphenicol (34 mg/ml) and incubated for 48h in 18°C with shaking 150 rpm. Bacteria were pelleted by centrifugation at 4500 rpm for 15 minutes at 4°C, frozen in liquid nitrogen and stored at −20°C. Pellet from 4 liters of culture was used for single purification. Pellet was resuspended in column buffer (50mM N2HPO4 pH 7.5, 150mM NaCl, 1mM EDTA) supplemented with 1 mM DTT, 1% Triton X-100, 1mM PMSF, a protein inhibitor cocktail (20 nM pepstatin; 6 nM leupeptin; 2 ng/ml chymostatin) and 50 µg/ml lysozyme, incubated for 20 minutes in 4 °C, then broken in a French pressure cell press MultisiFlex-C3 at 500 Bar. The homogenate was centrifuged in a Sorvall WX ULTRA SERIES ultracentrifuge, F37L rotor at 32000 rpm for 45 minutes at 4°C. The supernatant was loaded on 1 ml column with Glutathione Sepharose 4B resin, equilibrated by column buffer. ÄKTA Purifier system (GE Healthcare) was used for all purification steps. Unbound proteins were washed out with 20 CV of column buffer. GST-eIF4e was collected during 5 CV washing of elution buffer (50mM Na2HPO4 pH 8.5, 10mM L-Glutathione reduced, 1mM DTT). The elution fraction was mixed and diluted 3 times with water and loaded into Ion-exchange column (Resource S GE Healthcare), equilibrated by 50mM Na2HPO4 pH=8, 100 mM NaCl. GST-eIF4e was eluted by 0.1-1M NaCl gradient. All purification steps were analyzed by SDS-PAGE.

#### RNA isolation

Total RNA was isolated from cells with TRIzol reagent (Thermo Fisher Scientific) according to the manufacturer’s instructions, dissolved in nuclease-free water and stored at −20°C.

#### RT-qPCR

For the quantitative analysis, RNA was first treated with DNase (Thermo Fisher Scientific) for 30 min in 37°C and then reverse transcribed using SuperScript III (Thermo Fisher Scientific) and oligo(dT)20 and random-primers (Thermo Fisher Scientific). The quantitative PCR was performed with Platinum SYBR Green qPCR SuperMix-UDG (Thermo Fisher Scientific) using LightCycler 480 II (Roche) PCR device and appropriate primers listed in Supplementary Table 2. Gene expression for each sample was normalized to GAPDH. Differences were determined using the 2^−ΔΔC(t)^ calculation.

#### Northern Blotting

Northern blotting was performed as previously described (Mroczek et al., 2017). RNA samples were separated on 4% acrylamide gels containing 7M urea in 0.5x TBE buffer and transferred to a Hybond N+ membrane by electrotransfer in 0.5× TBE buffer. After transfer membranes, blots were stained with 0.03% methylene blue in 0.3 M NaAc pH 5.3 for 5 minutes at room temperature, scanned and then destained with water. RNA was immobilized on membranes by 254 nm UV light using a UVP CL-1000 crosslinker. Radioactive probes were labeled with a ^32^P (dATP) with a DECAprime II DNA Labeling Kit (Thermo Fisher Scientific). To obtain templates for probes labeling PCR on cDNA from B cells activated with LPS and IL4 for 7 days was conducted using primers listed in Supplementary Table 2.

Membranes were pre-hybridized in PerfectHyb Plus Hybridization Buffer (Sigma) for 1 hour 65°C and incubated with radioactive probes in PerfectHyb Plus Hybridization Buffer overnight 65°C. Then membranes were washed in 2xSSC with 0.1% SDS for 20 min, 0.5xSSC with 0.1% SDS for 20 min and 20min 0.1xSSC with 0.1% SDS for 20 min, scanned with Fuji Typhoon FLA 7000 (GE Healthcare Life Sciences) and analyzed with Multi Gauge software Ver. 2.0 (FUJI FILM).

#### mRNA enrichment with GST-eIF4E^K119A^ protein

Purification was performed as described previously with some modifications (Bajak and Hagedorn, 2008). Briefly, Glutathione-Sepharose 4B resin (GE Healthcare,) was incubated with GST-eIF4EK119A protein in sterile PBS (200 µl resin per 200 µg protein) for 1 hour at room temperature with rotation. Then the resin was washed 2 times with PBS and 3 times with buffer B (10 mM potassium phosphate buffer, pH 8.0, 100 mM KCl, 2 mM EDTA, 5% glycerol (Sigma), 0.005% Triton X-100 (Sigma), 6 mM DTT (A&A Biotechnology), and 20 U/mL Ribolock RNase Inhibitor (Thermo Scientific). 100 µg of total RNA, previously denaturated during 10 min in 70°C, was mixed with the prepared resin and incubated for 1 h at room temperature on an immunoprecipitation rotor. Then, the resin was washed 3 times with buffer B, 2 times with buffer B supplemented with 0.5 mM GDP (Sigma) and 2 times with buffer B without GDP. RNA was eluted from the resign by acid phenol:chloroform extraction and precipitated using 100% ethanol (Merck), 3M sodium acetate and GlycoBlue coprecipitant (Invitrogen).

#### Validation of *the* mRNA enrichment procedure with GST-eIF4E^K119A^ protein

150 µg of total RNA from HEK293T cells were subjected to mRNA enrichment with GST-eIF4E^K119A^ protein as described above. Then, both total and purified RNA were treated with DNase (Thermo Fisher Scientific) and 300 ng was used for reverse transcription (as described above). To estimate mRNA enrichment and rRNA removal efficiency cDNA was used for qPCR analysis, as described above.

#### RNAseq

All experiments were done in triplicate, where single replicate originated from TENT5C KO and WT littermates at age 12-15 weeks.

##### Cell culture and RNA retrieval

Naïve B cells were isolated and cultured as described above. Subsequently, after isolation cells were activated by the addition of 20 ng/ml IL-4 (Peprotech) and 20 μg/ml LPS (Santa Cruz) to the medium, followed by 7 days of incubation. Finally, RNA was isolated as described above.

##### Library preparation

Total RNA was treated with DNase (Invitrogen) for 30 min in 37°C and 1 µg of RNA was subjected to ribodepletion using a Ribo-Zero Kit (Illumina), according to the manufacturer’s recommendations and spiked-in with external RNA (ERCC RNA Spike-In Mix, Thermo Fisher Scientific). Strand-specific libraries were prepared using a dUTP protocol (KAPA Stranded RNA-Seq Library Preparation Kit), according to the manufacturer. Library quality was assessed using chip electrophoresis performed on an Agilent 2100 Bioanalyzer (Agilent Technologies, Inc.). The libraries were sequenced using an Illumina NextSeq500 sequencing platform to an average number of ∼1.5 × 10^7^ reads per library in the 75-nt paired-end mode.

#### Nanopore Direct RNA sequencing and polyadenylation analysis

##### RNA retrieval

Direct RNA sequencing was performed in duplicate, using the same input RNA (batches 1 and 2) as for the RNAseq experiment (described above). 100 µg of RNA was subjected to mRNA-enrichment with GST-eIF4E^K119A^ protein (as described above), followed by the ribodepletion of 2.5 µg RNA using a Ribo-Zero Kit (Illumina), according to the manufacturer’s recommendations.

##### Library preparation and sequencing

Nanopore direct RNA libraries were prepared from 500 ng of cap-enriched, rRNA-depleted mRNA with Direct RNA Sequencing Kit (ONT, SQK-RNA001). Instead of RNA CS, a 0.5 µl ERCC spike-in was added during RTA adapter ligation step and all remaining steps were performed according to the manufacturer’s instructions. Sequencing was performed with MinION device and Flow Cell (Type R9.4.1; RevC) and basecalled using Guppy 2.2.2 (Oxford Nanopore Technologies).

#### Co-Immunoprecipitation and Mass spectrometry analysis

Activated with LPS (20 μg/ml) and IL4 (20 ng/ml) TENT5C-GFP B cells were crosslinked with 1 mM DSP (dithiobis(succinimidyl propionate); Invitrogen) for 1 h before stopping the reaction with 50 mM Tris pH 8.0. After washing with PBS-supplemented 50 mM Tris pH 8.0, cells will be flash-frozen in liquid nitrogen, thawed on ice, and incubated for 30 min at 4°C with gentle rotation in 3 mL LB buffer (50 mM Tris, 150 mM NaCl, 0.5% Triton-X100, 1 mM DTT, supplemented with proteases and phosphatase inhibitors; Invitrogen). Next, the lysates were sonicated for 30 min with a Bioruptor Plus (Diagenode), followed by clarification by centrifugation. Immunoprecipitations were performed using a GFP-Trap (Chromotek). After 2 h of incubation, the beads were washed 6 times with LB buffer and finally, the proteins were eluted with 50 mM glycine pH 2.8. After neutralization with Tris pH 8.0, proteins were precipitated with PRM reagent (0.05 mM pyrogallol red, 0.16 mM sodium molybdate, 1 mM sodium oxalate, 50 mM succinic acid; pH 2.5 (Sigma-Aldrich)) prior to MS analysis in the Laboratory of Mass Spectrometry, IBB PAS (Marshall et al., 1995).

#### Western Blotting

For western blot analysis equal amount of cells were lysed with 0.1% NP40 in PBS supplemented with protease inhibitors and viscolase (A&A Biotechnology) for 30 min in 37°C with shaking 600 rpm, then Laemmli buffer was added and samples were denaturated for 10 min in 100°C. For secreted proteins analysis, the cell culture medium was collected and centrifuged twice 15 min 13 500 rpm, then the supernatant was collected, Laemmli buffer was added and samples were denaturated for 10 min in 100°C. Samples were separated on 12-15% SDS-PAGE gels, proteins were transferred to Protran nitrocellulose membranes (GE Healthcare) and then membranes were stained with 0.3% w/v Ponceau S in 3% v/v acetic acid and digitized. Membranes were incubated with 5% milk or 5% BSA in TBST buffer according to the technical recommendations of the antibodies’ suppliers for 1 hour followed by incubation with specific primary antibodies (listed in the Key Resources Table) diluted 1:10 000 (α-tubulin), 1:5 000 (IgG, Igλ, IgGκ), 1:3 000 (IL6, GRP94), 1:2 000 (PERK, Xbp1 or 1:1000 (Ire1, ATF6, FLAG) overnight in 4°C. Membranes were washed 3 times in TBST buffer, incubated with HRP-conjugated secondary antibodies (anti-mouse diluted 1:5 000 anti-rabbit diluted 1:3 000) for 2 hours at RT. Membranes were washed 3 times in TBST buffer and proteins were visualized by enhanced chemiluminescence acquired on X-ray film.

#### Flow Cytometry Analysis

Splenocytes and bone marrow were isolated as described above, and after depletion of red blood cells with ACK buffer cells were stained respectively. Designed staining panels were based on the „Flow cytometry tools for the study of B cell biology” (BD Pharmingen) (Pracht et al., 2017). The antibodies and other reagents used for flow cytometry analysis are listed in the Key Resources Table. Samples were measured with BD LSRFortessa™ (BD) and analyzed using FlowJo (Data Analysis Software v10).

#### Surface staining for analysis of early developmental stages of B cells

1.5×10^6^ cells isolated from bone marrow or spleen were pelleted and incubated with Fc Block (anti-CD16/32) for 10 min RT. After washing with FACS buffer (0.2% BSA in PBS) cells isolated from bone marrow were stained with anti-CD19 BUV395, anti-CD43 BV421, anti-CD23 BV421, anti-IgM FITC, anti-CD249 PE, anti-CD93 PE, anti-IgD BV605, anti-CD45R/B220 APC, anti-CD24 PerCP Cy5.5, anti-CD21 PerCP Cy5.5 antibodies for 30 min in 4°C, protected from light. In case, when two antibodies produced with BD Horizon technology were used in one staining panel, Brilliant Stain Buffer was used to prepare the antibody mix. After staining, cells were washed with FACS buffer and then stained with LIVE/DEAD™ Fixable IR Dead Cell Stain Kit for 20 min in 4°C, protected from light. Finally, cells were washed and analyzed in terms of calculation PreProB, ProB, PreB, immature, transitional B, early and late mature B, T1, T2, T3, follicular B I and II and marginal cell subsets (Marginal Zone and Marginal Zone Progenitor).

#### Surface staining for analysis of plasmablasts and plasma cells

2.5×10^6^ cells isolated from bone marrow or spleen were incubated with Fc Block (anti-CD16/32) for 10 min RT, washed with FACS buffer and incubated with anti-CD19 BUV395, anti-CD19 AF700, anti-IgM FITC, anti-IgA PE, anti-CD138 BV605, anti-CD138 PE, anti-IgG1 AF647, anti-CD45R/B220 PerCP Cy5.5 antibodies for 30 min in 4°C, protected from light. In case, when two antibodies produced with BD Horizon technology were used in one staining panel, Brilliant Stain Buffer was used to prepare the antibody mix. Next cells were washed with FACS buffer and then stained with LIVE/DEAD™ Fixable Near-IR or Violet Dead Cell Stain Kit for 20 min in 4°C, protected from light. Finally, cells were washed and analyzed in terms of calculation dividing plasmablasts, early plasmocytes and mature resting plasmocytes.

#### Cytoplasmic staining for analysis of intracellular immunoglobulins

2.5×10^6^ cells isolated from bone marrow or spleen were incubated with Fc Block (anti-CD16/32) for 10 min RT. After washing with FACS buffer (0.2% BSA in PBS) cells isolated from bone marrow were stained with anti-CD19 BUV395, anti-CD138 BV605, anti-CD45R/B220 PerCP Cy5.5 antibodies for 30 min in 4°C, protected from light. Cells were washed with FACS buffer and then incubated in Fixation/Permeabilization Solution for 20 min in 4°C. Upon washing in Wash/Permeabilization solution, cells were stained with anti-IgM FITC, anti-IgA PE, anti-IgG1 AF647 for 30 min in 4°C. After washing cells were analyzed in terms of calculation intracellular/cytoplasmic intracellular immunoglobulin content.

#### Intracellular ER staining with for live-cell imaging

1×10^6^ cells were resuspended in HSBB buffer and then were incubated with 0.75 µl of ER-Tracker™ Red (BODIPY™ TR Glibenclamide) for 30 min at 4°C, in the dark.

#### Splenocyte fractionation

Activated B cells isolated from TENT5C-FLAG mice were fractionated using a Subcellular Protein Fractionation Kit for Cultured Cells (Invitrogen) according to the manufacturer’s instruction. The resulting fractions were analyzed by western blot.

#### ER stress analysis

WT and TENT5C KO B cells were activated with LPS (20 μg/ml) and IL-4 (20 ng/ml) for 3 days. Then tunicamycin (Sigma) was added to a final concentration 1 or 2.5 µg/ml for 5 hours and cells were collected. RNA was isolated using TRI reagent (Sigma) according to the manufacturer’s instructions and estimations of Xbp1 splicing and expression level of PERK, CHOP, GRP94, Ero1lB and Ire1α genes were carried out by RT-qPCR reactions as described above. Cells were also used for protein isolation and western blot analysis of UPR markers level (XBP1, Ire1, PERK, ATF6) as described above.

### QUANTIFICATION AND STATISTICAL ANALYSIS

Statistical analysis of quantitative data was performed using Prism software (GraphPad), unless otherwise stated. The statistical tests used in each instance are mentioned in the figure legends. All data were checked for normality using Shapiro-Wilk test. Outliers were identified in GraphPad Prism using ROUT method, with Q (FDR) value set to 1%, and removed from subsequent analyses.

Data are presented as scatter dot plots or bar plots, with mean values indicated and standard deviation shown as the error bars. Statistical significance is marked as: ns - not significant, ∗ p < 0.05, ∗∗ p < 0.01, ∗∗∗ p < 0.001, ∗∗∗∗ p < 0.0001.

#### Differential Expression Analysis

Obtained sequencing reads were quality filtered using cutadapt (v. 1.18), to remove adapter sequences, low-quality fragments (minimum quality score was set to 20) and too short sequences (threshold set to 30 nt) (Martin, 2011). Such quality filtered reads were aligned to the mouse genome (GRCm38, downloaded from the ENSEMBL FTP site, release 94) using the STAR split read aligner (v. 2.6.1a) (Dobin et al., 2013). Counts for each transcript were collected using featureCounts from Subread package (v. 1.6.3), with options -Q 10 -p -B -C -s 2 -g gene_id -t exon and Gencode vM19 annotation (Liao et al., 2014). Statistical analysis of differential expression was performed using the DESeq2 (v. 1.22) Bioconductor package (Love et al., 2014), using default settings, correcting for the batch effect.

#### Polyadenylation analysis using Nanopore Direct RNA Sequencing

To generate B cells specific reference transcriptome we used FLAIR v1.2 (https://github.com/BrooksLabUCSC/flair) (Tang A et al., 2018). In detail, nanopore reads from all analyzed samples were mapped to the GRCm38 genome with flair.py align. Misaligned splice sites were corrected with flair correct and splice junctions file created by STAR mapping of all Illumina reads to GRCm38 genome. High confidence isoforms dataset was created using flair collapse, and further annotated against GencodeVM19 using spliced_bam2gff from pinfish (https://github.com/nanoporetech/pinfish) and GFFCompare (v. 0.10.6, https://ccb.jhu.edu/software/stringtie/gffcompare.shtml).

Nanopore reads were mapped back to such created reference transcriptome using Minimap 2.14 (Li, 2018). The poly(A) tail lengths for each read were estimated using Nanopolish 0.10.2 polya function (Workman et al., 2018). In subsequent analyses, only length estimates with QC tag reported by Nanopolish as PASS were considered. Statistical analysis was performed using functions provided in the NanoTail R package (https://github.com/smaegol/nanotail). In detail, we used the Generalized Linear Model approach, with log2(polya length) as a response variable. To correct for the batch effect, a replicate identifier was used as one of the predictors, in addition to condition (TENT5CKO/WT) identifier. Collected P values (for the condition effect) were adjusted for multiple comparisons using the Benjamini-Hochberg method. Transcripts were considered as having poly(A) tail significantly changed between analyzed conditions, if the adjusted p.value was less than 0.05, the absolute difference in the median tail length was at least 10, and there were at least 10 supporting reads for each condition.

### DATA AND CODE AVAILABILITY

All mRNA expression data that support the findings of this study have been deposited at GEO under accession number GSE132883 (temporarily token for a read-only access for reviewers: qruhwomczzkjfwj). The raw basecalled data from Nanopore Direct RNA Sequencing are deposited at European Nucleotide Archive, under study accession number PRJEB33089.

---

**Supplementary Figure 1.**
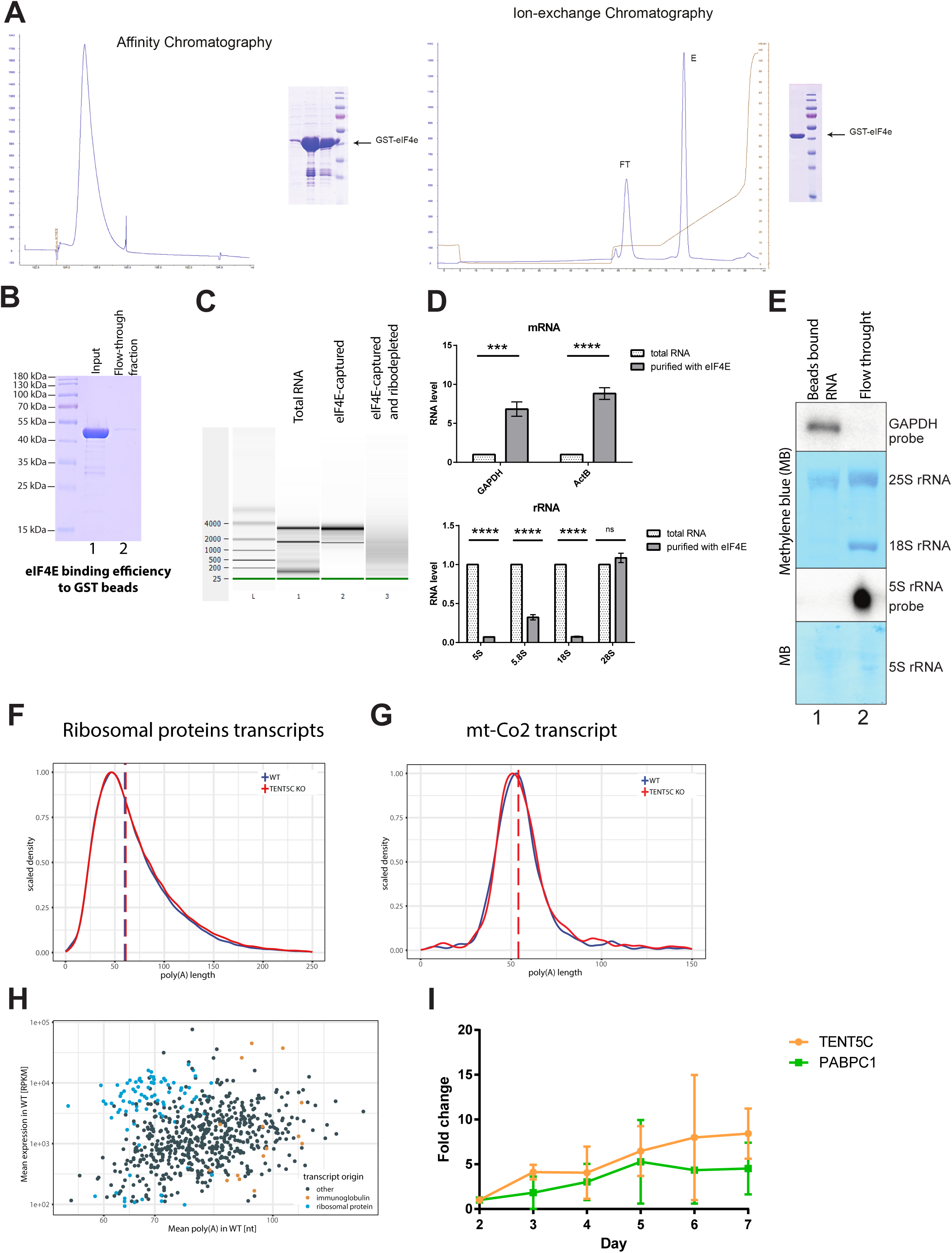
Modified direct RNA Sequencing is reliable method to measure poly(A) tails in responding B cells. (Related to Figure 1). (A) Purification of the GST-eIF4E protein. Affinity chromatography and ion-exchange chromatography profiles with SDS-PAGE analysis of the purified GST-eIF4E protein. (B) Validation of the purified eIF4E batch binding to GST beads. Coomassie Blue-stained SDS-PAGE gel analysis of purified eIF4E input (lane 1) and flow-through fraction (lane 2) after incubation with GST beads. (C) Agilent 2100 Bioanalyzer electropherogram profiles of total RNA, eIF4E-captured RNA and both eIF4E-captured and ribodepleted RNA from WT B cells activated with LPS and IL-4 for 7 days. (D) The efficiency of mRNA enrichment (higher panel) and rRNA removal (lower panel) after eIF4E-capture assessed by qPCR analysis. Total RNA form HEK293T cells were purified with eIF4E protein. Then input and purified RNA were subjected to qPCR. Bars represents mean values of technical repeats ±SD (n=3), P values were calculated using unpaired Student’s t-test. (E) Northern blot analysis of eIF4E-capture efficiency. Total RNA form HEK293T cells were purified with eIF4E protein. Then purified RNA and unbound flow-through fraction were compared. (F) (G) Nanopore-based poly(A) lengths profiling of B cells isolated from WT and TENT5C KO, activated with LPS and IL4 for 7 days. Shown are density distribution plots, scaled to a maximum of 1, for: (F) ribosomal proteins transcripts, (G) mitochondrially encoded cytochrome c oxidase II transcript. Vertical dashed lines represent median poly(A) lengths for each condition (H) Scatter plot showing the relation between mean poly(A) length (x-axis, obtained with Nanopore direct RNA sequencing for WT) and expression (y-axis, obtained with Illumina RNAseq), for each individual transcript. Immunoglobulins and highly expressed ribosomal proteins transcripts are marked in orange and blue, respectively. (I) qPCR analysis of TENT5C and PABPC1 expression levels in WT B cells (n=4) activated with LPS and IL4 for 2-7 days. Pearson’s correlation between TENT5C and PABPC1 expression r=0.88.

**Supplementary Figure 2.**
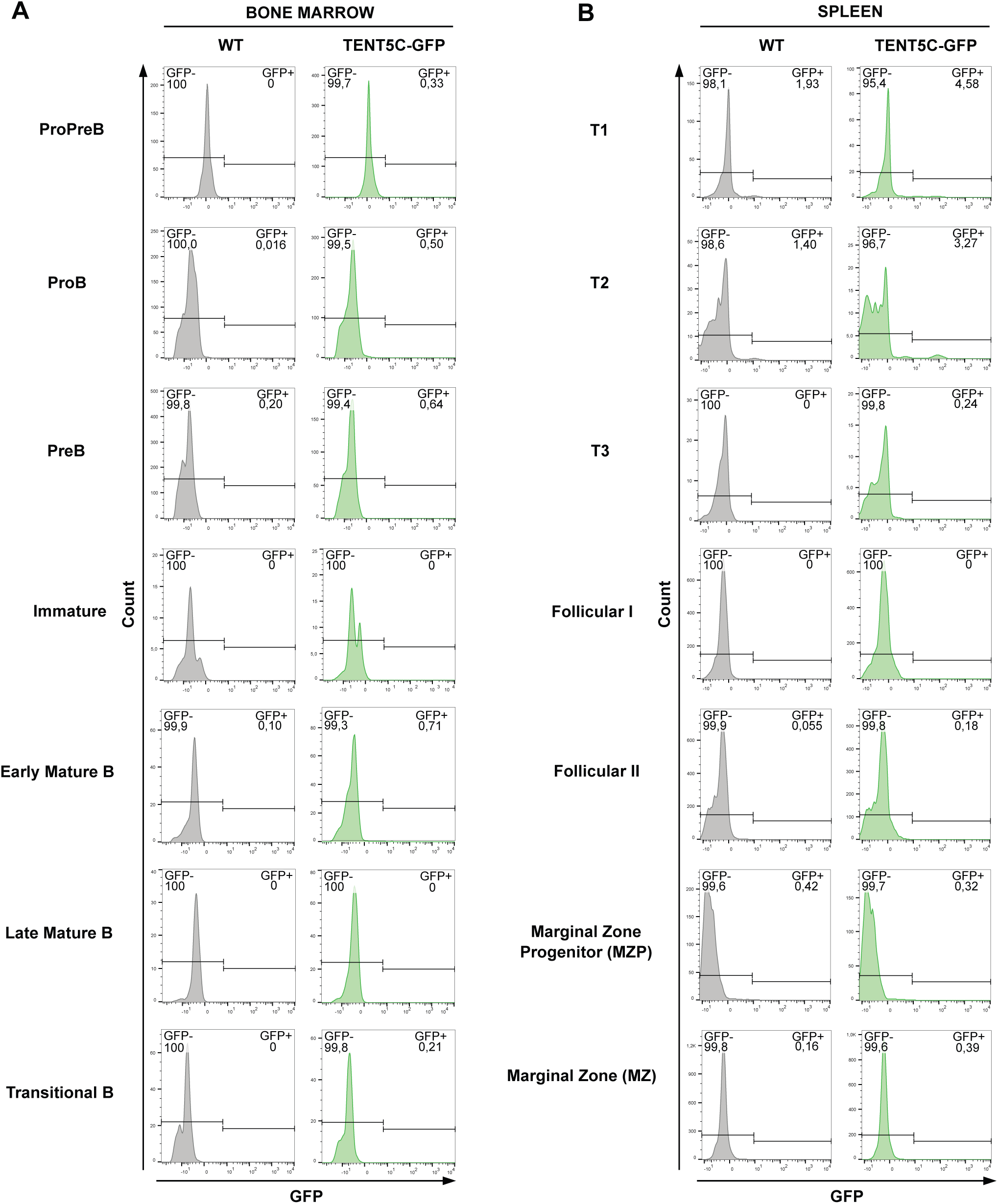
Flow cytometry analysis of TENT5C-GFP expression in B cells subpopulations. (Related to Figure 3). (A, B) Histograms showing GFP fluorescence intensity in different B lymphocytes subsets from WT or TENT5C-GFP bone marrow (A) and spleen (B). Cells from bone marrow were stained based on the CD19, CD43, CD45R, CD24, CD249, IgM and IgD markers. Cells from spleen were stained based on the CD19, CD23, CD93, CD45R, CD21, IgM and IgD markers. Staining panels distinguish following B cell subsets: ProPreB, ProB, PreB, Immature, Early Mature B, Late Mature B, Transitional B, T1/T2/T3, Follicular I and II, Marginal Zone Progenitors and Marginal Zone. See also gating strategy in Supplementary Figure 3.

**Supplementary Figure 3.**
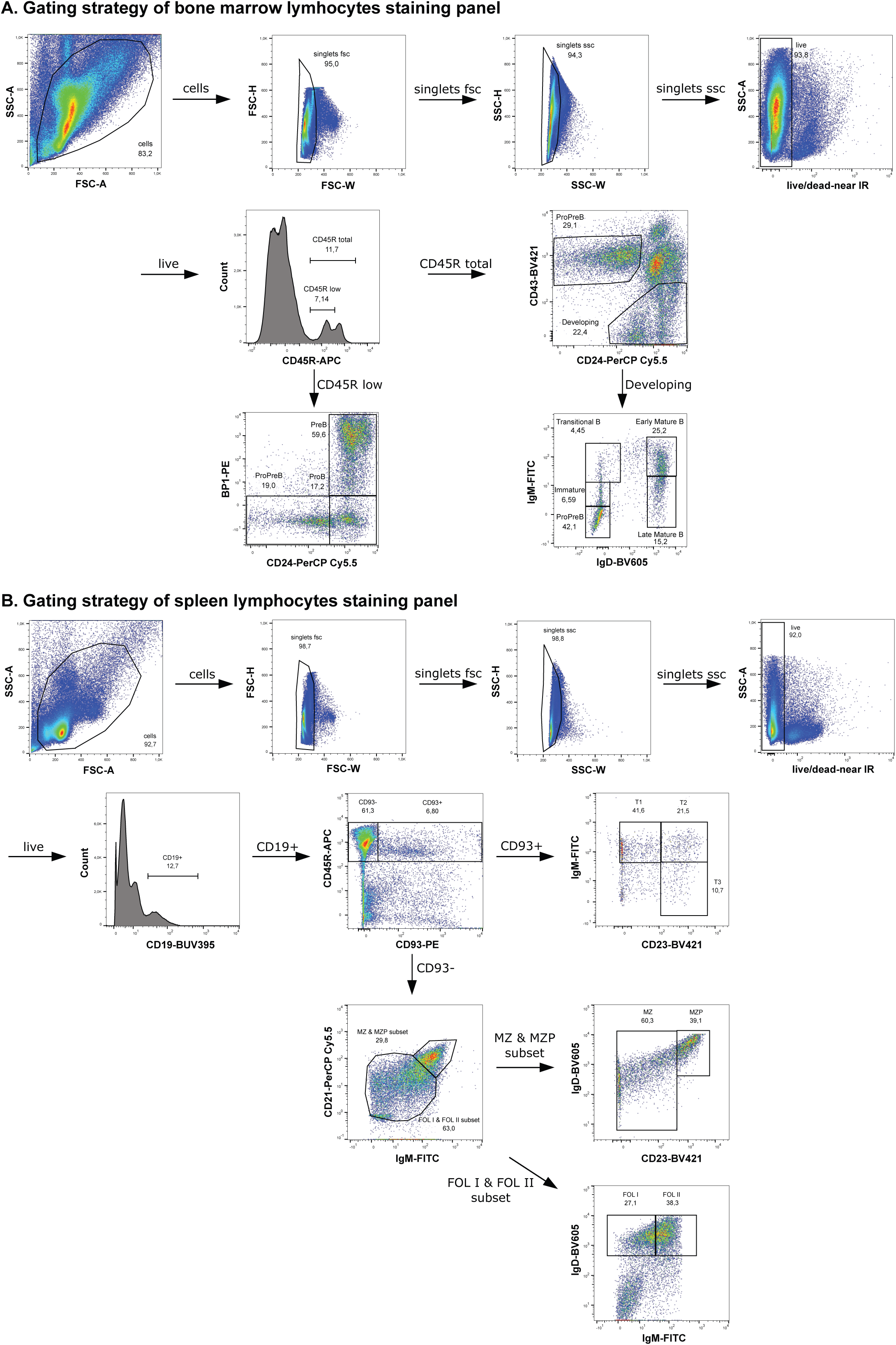
Gating strategy of bone marrow B cells staining panel (A) and spleen B cells staining panel (B). (Related to Supplementary Figure 2 and Supplementary Figure 6). (A) The 8-colour flow cytometry analysis of B cells isolated from bone marrow. Cells were stained with fluorochrome-conjugated antibodies against: CD19-BUV395, CD43-BV421, IgM-FITC, CD249-PE, IgD-BV605, CD45R-APC, CD24-PerCP Cy5.5. When TENT5C-GFP level was checked antibody against IgM-BV510 was used (see Supplementary Figure 2). To exclude dead cells LIVE/DEAD™ Fixable Near-IR Dead Cell Stain Kit was used. (B) A 8-colour flow cytometry analysis of B cells isolated from spleen. Cells were stained with fluorochrome-conjugated antibodies against: CD19-BUV395, CD23-BV421, IgM-FITC, CD93-PE, IgD-BV605, CD45R-APC, CD21-PerCP Cy5.5 When TENT5C-GFP level was checked antibody against IgM-BV510 was used (see Supplementary Figure 2). To exclude dead cells LIVE/DEAD™ Fixable Near-IR Dead Cell Stain Kit was used.

**Supplementary Figure 4.**
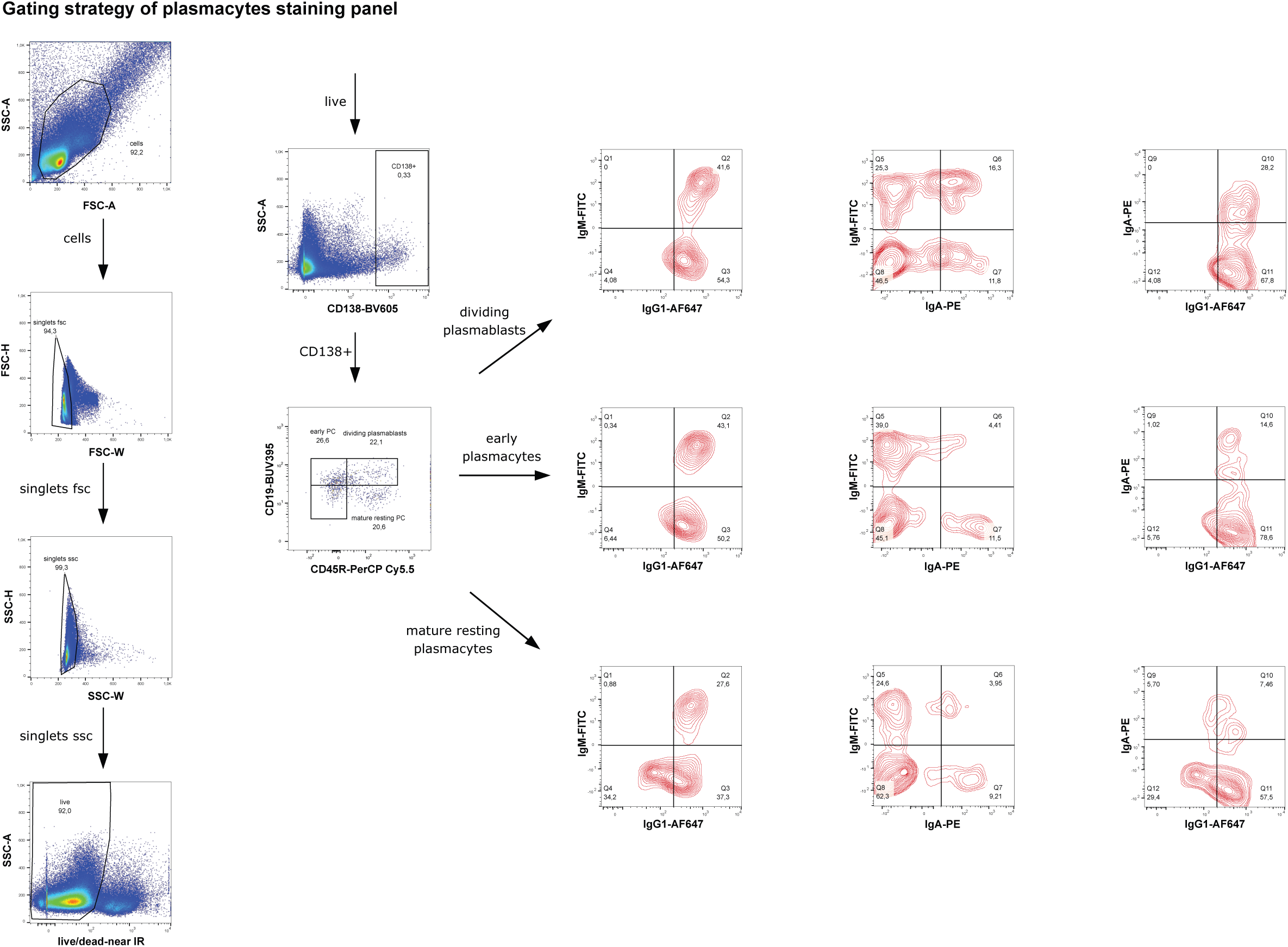
Gating strategy of plasmacytes staining panel. (Related to Figures 2, 3, 5, 6 and 7). A 7-colour flow cytometry analysis of plasma cells isolated from bone marrow or spleen. Cells were stained with fluorochrome-conjugated antibodies against: CD19-BUV395, IgM-FITC, IgA-PE, CD138-BV605, IgG1-AF647, CD45R-PerCP Cy5.5 (see analysis on Figure 5, 6 and 7). For intracellular (cytoplasmic) immunoglobulins measurement (see Figure 2) the same gating strategy was applied. When TENT5C-GFP level was analyzed antibody against CD138-PE and CD19-AF700 was used (see Figure 3), for measuring ER additionally, the ER-tracker-Red was used (Figure 7). To exclude dead cells LIVE/DEAD™ Fixable Near-IR or Violet Dead Cell Stain Kit was used.

**Supplementary Figure 5.**
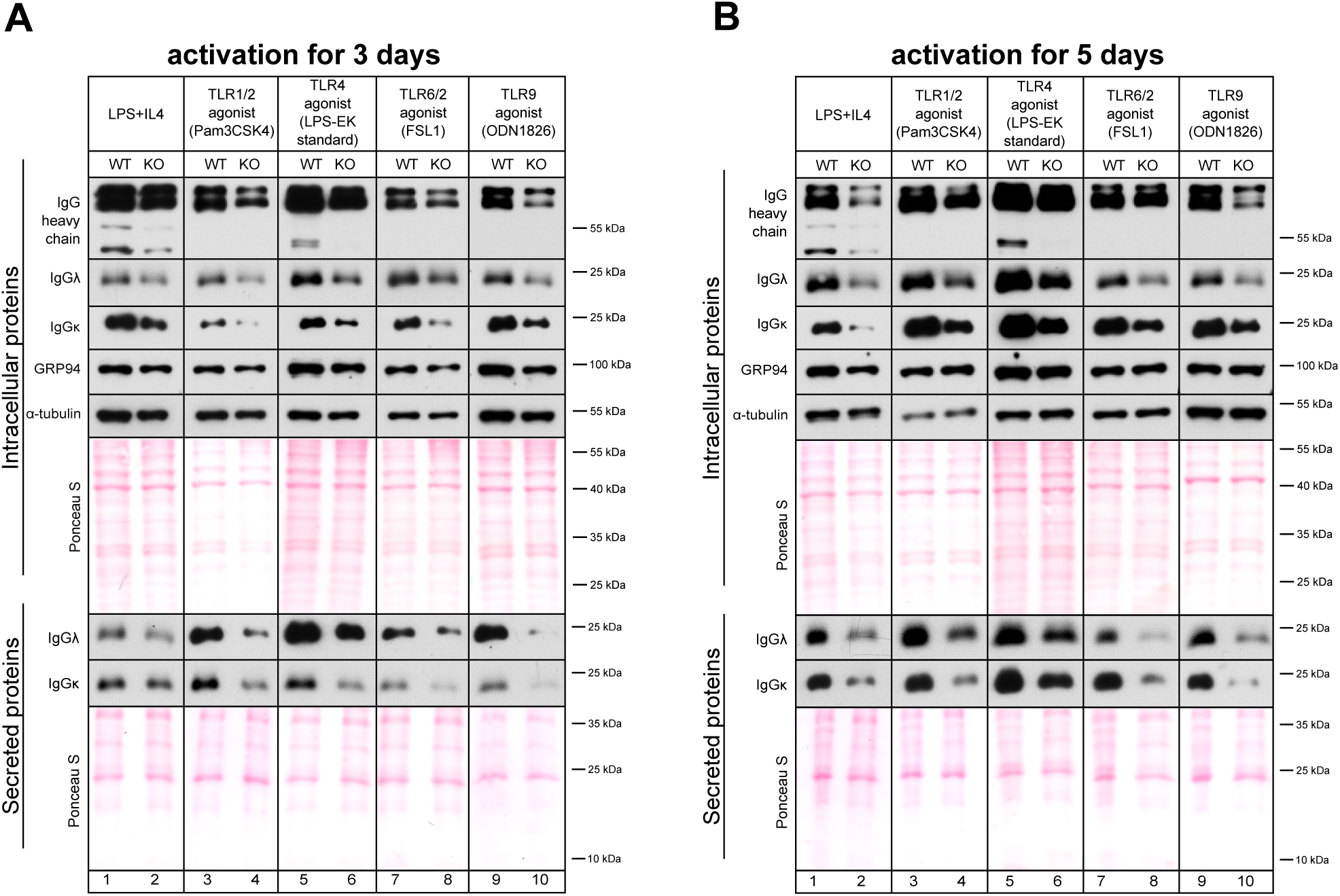
TENT5C KO B cells activated by innate signaling pathways produce less antibodies. (Related to Figure 4) **(A, B)** Western blot analysis of IgG heavy and light chains (λ and κ), intracellular and secreted level, in TENT5C KO and WT B cells, activated with TLR agonists: LPS/ IL4 (positive control; lanes 1,2), TLR 1/2 (lanes 3,4), TLR4 (lanes 5,6), TLR6/2 (lanes 7,8) and TLR9 (lanes 9,10) for 3 (A) or 5 (B) days. GRP94 was used as an activation marker, α-tubulin and Ponceau S staining were used as loading controls.

**Supplementary Figure 6.**
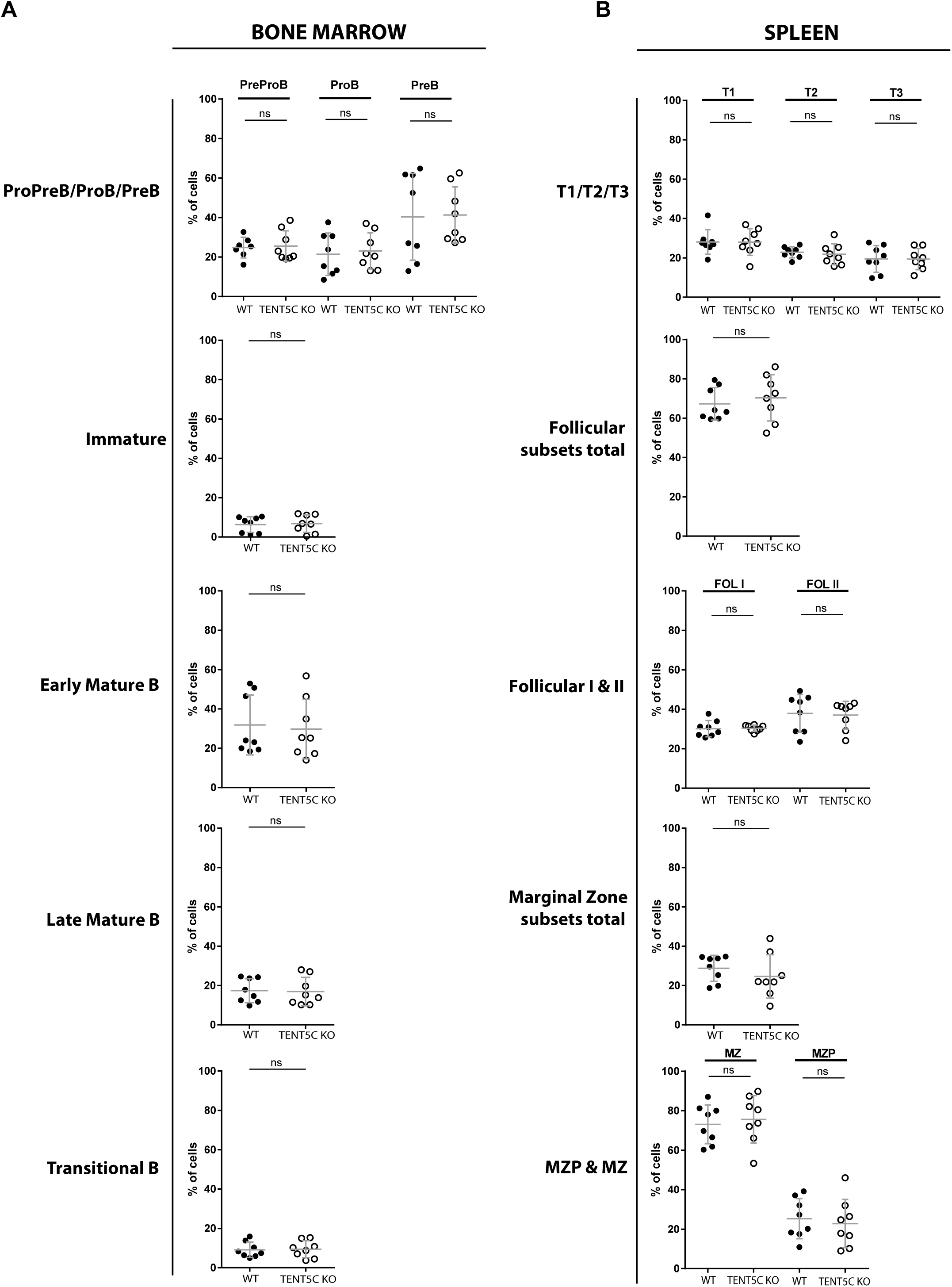
TENT5C expression does not influence the early stages of B cells development (related to Figure 5). (A, B) Comparison of different B cells subsets isolated from WT or TENT5C KO from bone marrow (A) and spleen (B) based on the flow cytometry results. Subpopulations of ProPreB, ProB, ProB, Immature, Early Mature B, Late Mature B, Transitional B, T1/T2/T3, Follicular I & II, Marginal Zone Progenitors (MZP) and Marginal Zone (MZ) were checked based on CD19, CD43, CD45R, CD24, CD249, IgM, IgD, CD23, CD93, CD21 markers. See gating strategy in Supplementary Figure 3. Data are shown as a mean of WT n=8, TENT5C KO n=8 mice, two-way ANOVA with Bonferroni post hoc test.

## Literature.

Alasaad, S., Rossi, L., Maione, S., Sartore, S., Soriguer, R.C., Perez, J.M., Rasero, R., Zhu, X.Q., and Soglia, D. (2008). HotSHOT Plus ThermalSHOCK, a new and efficient technique for preparation of PCR-quality mite genomic DNA. Parasitol Res 103, 1455–1457.

Amend, S.R., Valkenburg, K.C., and Pienta, K.J. (2016). Murine Hind Limb Long Bone Dissection and Bone Marrow Isolation. J Vis Exp.

Aragon I.V., Barrington R.A., Jackowski S., Mori K., and Brewer J.W. (2012). The specialized unfolded protein response of B lymphocytes: ATF6α-independent development of antibody-secreting B cells. Mol Immunol. 51, 347–55.

Bajak, E.Z., and Hagedorn, C.H. (2008). Efficient 5’ cap-dependent RNA purification: use in identifying and studying subsets of RNA. Methods in molecular biology 419, 147–160.

Braakman, I., and Hebert, D.N. (2013). Protein folding in the endoplasmic reticulum. Cold Spring Harbor perspectives in biology 5, a013201.

Chang, H., Lim, J., Ha, M., and Kim, V.N. (2014). TAIL-seq: genome-wide determination of poly(A) tail length and 3’ end modifications. Molecular cell 53, 1044–1052.

Choi, Y.H., and Hagedorn, C.H. (2003). Purifying mRNAs with a high-affinity eIF4E mutant identifies the short 3’ poly(A) end phenotype. Proceedings of the National Academy of Sciences of the United States of America 100, 7033–7038.

Chorghade S., Seimetz J., Emmons R., Yang J., Bresson S.M., Lisio M., Parise G., Conrad N.K., and Kalsotra A. (2017). Poly(A) tail length regulates PABPC1 expression to tune translation in the heart. Elife. 6, e24139.

Danger, R., Braza, F., Giral, M., Soulillou, J.P., and Brouard, S. (2014). MicroRNAs, Major Players in B Cells Homeostasis and Function. Front Immunol 5, 98.

De Silva, N.S., and Klein, U. (2015). Dynamics of B cells in germinal centres. Nat Rev Immunol 15, 137–148.

Diaz-Munoz, M.D., Monzon-Casanova, E., and Turner, M. (2017). Characterization of the B Cell Transcriptome Bound by RNA-Binding Proteins with iCLIP. Methods in molecular biology 1623, 159–179.

Dobin, A., Davis, C.A., Schlesinger, F., Drenkow, J., Zaleski, C., Jha, S., Batut, P., Chaisson, M., and Gingeras, T.R. (2013). STAR: ultrafast universal RNA-seq aligner. Bioinformatics 29, 15–21.

Eaton J.D., Davidson L., Bauer D.L.V., Natsume T., Kanemaki M.T., and West S. (2018). Xrn2 accelerates termination by RNA polymerase II, which is underpinned by CPSF73 activity. Genes Dev. 32, 127–139.

Enders, A., Short, A., Miosge, L.A., Bergmann, H., Sontani, Y., Bertram, E.M., Whittle, B., Balakishnan, B., Yoshida, K., Sjollema, G., et al. (2014). Zinc-finger protein ZFP318 is essential for expression of IgD, the alternatively spliced Igh product made by mature B lymphocytes. Proceedings of the National Academy of Sciences of the United States of America 111, 4513–4518.

Feng, Y., Zhang, Y., Ying, C., Wang, D., and Du, C. (2015). Nanopore-based fourth-generation DNA sequencing technology. Genomics Proteomics Bioinformatics 13, 4–16.

Friday, A.J., and Keiper, B.D. (2015). Positive mRNA Translational Control in Germ Cells by Initiation Factor Selectivity. Biomed Res Int 2015, 327963.

Garalde, D.R., Snell, E.A., Jachimowicz, D., Sipos, B., Lloyd, J.H., Bruce, M., Pantic, N., Admassu, T., James, P., Warland, A., et al. (2018). Highly parallel direct RNA sequencing on an array of nanopores. Nature methods 15, 201–206.

Gass, J.N., Gunn, K.E., Sriburi, R., and Brewer, J.W. (2004). Stressed-out B cells? Plasma-cell differentiation and the unfolded protein response. Trends Immunol 25, 17–24.

Goldfinger, M., Shmuel, M., Benhamron, S., and Tirosh, B. (2011). Protein synthesis in plasma cells is regulated by crosstalk between endoplasmic reticulum stress and mTOR signaling. Eur J Immunol 41, 491–502.

Harrison, P.F., Powell, D.R., Clancy, J.L., Preiss, T., Boag, P.R., Traven, A., Seemann, T., and Beilharz, T.H. (2015). PAT-seq: a method to study the integration of 3’-UTR dynamics with gene expression in the eukaryotic transcriptome. Rna 21, 1502–1510.

Houseley, J., and Tollervey, D. (2009). The many pathways of RNA degradation. Cell 136, 763–776.

Hrit, J., Raynard, N., Van Etten, J., Sankar, K., Petterson, A., and Goldstrohm, A.C. (2014). In vitro analysis of RNA degradation catalyzed by deadenylase enzymes. Methods in molecular biology 1125, 325–339.

Kakiuchi-Kiyota S., Arnold L.L., Yokohira M., Koza-Taylor P., Suzuki S., Varney M., Pennington K.L., and Cohen S.M. (2011). Evaluation of direct and indirect effects of the PPARγ agonist troglitazone on mouse endothelial cell proliferation. Toxicol Pathol. 39, 1032–45.

Kang, S.M., Yoo, D.G., Kim, M.C., Song, J.M., Park, M.K., O, E., Quan, F.S., Akira, S., and Compans, R.W. (2011). MyD88 plays an essential role in inducing B cells capable of differentiating into antibody-secreting cells after vaccination. J Virol 85, 11391–11400.

Kirk, S.J., Cliff, J.M., Thomas, J.A., and Ward, T.H. (2010). Biogenesis of secretory organelles during B cell differentiation. J Leukoc Biol 87, 245–255.

Kobyłecki K., Drążkowska K., Kuliński T.M., Dziembowski A., and Tomecki R. (2018). Elimination of 01/A’-A0 pre-rRNA processing by-product in human cells involves cooperative action of two nuclear exosome-associated nucleases: RRP6 and DIS3. RNA. 24, 1677–1692.

Kolesnikova I.S., Dolskiya A.A., Lemskaya N.A., Maksimova Y.V., Shorina A.R., Graphodatsky A.S., Galanina E.M., and Yudkin D.V. (2018). Alteration of rRNA gene copy number and expression in patients with intellectual disability and heteromorphic acrocentric chromosomes. Egyptian Journal of Medical Human Genetics. 19, 129–134.

Koralov, S.B., Muljo, S.A., Galler, G.R., Krek, A., Chakraborty, T., Kanellopoulou, C., Jensen, K., Cobb, B.S., Merkenschlager, M., Rajewsky, N., and Rajewsky, K. (2008). Dicer ablation affects antibody diversity and cell survival in the B lymphocyte lineage. Cell 132, 860–874.

Kuchta, K., Knizewski, L., Wyrwicz, L.S., Rychlewski, L., and Ginalski, K. (2009). Comprehensive classification of nucleotidyltransferase fold proteins: identification of novel families and their representatives in human. Nucleic acids research 37, 7701–7714.

Kuchta, K., Muszewska, A., Knizewski, L., Steczkiewicz, K., Wyrwicz, L.S., Pawlowski, K., Rychlewski, L., and Ginalski, K. (2016). FAM46 proteins are novel eukaryotic non-canonical poly(A) polymerases. Nucleic acids research 44, 3534–3548.

Legnini I., A.J., Karaiskos N., Ayoub S., Rajewsky N. (2018). Full-length mRNA sequencing reveals principles of poly(A) tail length control. BiorxiV, https://doi.org/10.1101/547034.

Li, H. (2018). Minimap2: pairwise alignment for nucleotide sequences. Bioinformatics 34, 3094–3100.

Li, M.Z., and Elledge, S.J. (2012). SLIC: a method for sequence- and ligation-independent cloning. Methods in molecular biology 852, 51–59.

Liao, Y., Smyth, G.K., and Shi, W. (2014). featureCounts: an efficient general purpose program for assigning sequence reads to genomic features. Bioinformatics 30, 923–930.

Lima, S.A., Chipman, L.B., Nicholson, A.L., Chen, Y.H., Yee, B.A., Yeo, G.W., Coller, J., and Pasquinelli, A.E. (2017). Short poly(A) tails are a conserved feature of highly expressed genes. Nature structural & molecular biology 24, 1057–1063.

Liu, B., and Li, Z. (2008). Endoplasmic reticulum HSP90b1 (gp96, grp94) optimizes B-cell function via chaperoning integrin and TLR but not immunoglobulin. Blood 112, 1223–1230.

Love, M.I., Huber, W., and Anders, S. (2014). Moderated estimation of fold change and dispersion for RNA-seq data with DESeq2. Genome biology 15, 550.

Lynes, E.M., and Simmen, T. (2011). Urban planning of the endoplasmic reticulum (ER): how diverse mechanisms segregate the many functions of the ER. Biochimica et biophysica acta 1813, 1893–1905.

Marshall, T., Abbott, N.J., Fox, P., and Williams, K.M. (1995). Protein concentration by precipitation with pyrogallol red prior to electrophoresis. Electrophoresis 16, 28–31.

Martin, M. (2011). Cutadapt removes adapter sequences from high-throughput sequencing reads. EMBnet.Journal 17, 10–12.

Mroczek, S., Chlebowska, J., Kulinski, T.M., Gewartowska, O., Gruchota, J., Cysewski, D., Liudkovska, V., Borsuk, E., Nowis, D., and Dziembowski, A. (2017). The non-canonical poly(A) polymerase FAM46C acts as an onco-suppressor in multiple myeloma. Nat Commun 8, 619.

Nicholson, A.L., and Pasquinelli, A.E. (2018). Tales of Detailed Poly(A) Tails. Trends in cell biology.

Pasare, C., and Medzhitov, R. (2005). Control of B-cell responses by Toll-like receptors. Nature 438, 364–368.

Oslowski C.M., and Urano F. (2011). Measuring ER stress and the unfolded protein response using mammalian tissue culture system. Methods Enzymol. 490, 71–92.

Peng, Y., Yuan, J., Zhang, Z., and Chang, X. (2017). Cytoplasmic poly(A)-binding protein 1 (PABPC1) interacts with the RNA-binding protein hnRNPLL and thereby regulates immunoglobulin secretion in plasma cells. The Journal of biological chemistry 292, 12285–12295.

Pioli, P.D., Debnath, I., Weis, J.J., and Weis, J.H. (2014). Zfp318 regulates IgD expression by abrogating transcription termination within the Ighm/Ighd locus. J Immunol 193, 2546–2553.

Pracht, K., Meinzinger, J., Daum, P., Schulz, S.R., Reimer, D., Hauke, M., Roth, E., Mielenz, D., Berek, C., Corte-Real, J., et al. (2017). A new staining protocol for detection of murine antibody-secreting plasma cell subsets by flow cytometry. Eur J Immunol 47, 1389–1392.

Reimold, A.M., Iwakoshi, N.N., Manis, J., Vallabhajosyula, P., Szomolanyi-Tsuda, E., Gravallese, E.M., Friend, D., Grusby, M.J., Alt, F., and Glimcher, L.H. (2001). Plasma cell differentiation requires the transcription factor XBP-1. Nature 412, 300–307.

Schuck, S., Prinz, W.A., Thorn, K.S., Voss, C., and Walter, P. (2009). Membrane expansion alleviates endoplasmic reticulum stress independently of the unfolded protein response. The Journal of cell biology 187, 525–536.

Serrano R.L., Yu W., and Terkeltaub R. (2014). Mono-allelic and bi-allelic ENPP1 deficiency promote post-injury neointimal hyperplasia associated with increased C/EBP homologous protein expression. Atherosclerosis. 233, 493–502.

Shi, W., Liao, Y., Willis, S.N., Taubenheim, N., Inouye, M., Tarlinton, D.M., Smyth, G.K., Hodgkin, P.D., Nutt, S.L., and Corcoran, L.M. (2015). Transcriptional profiling of mouse B cell terminal differentiation defines a signature for antibody-secreting plasma cells. Nat Immunol 16, 663–673.

Siwaszek, A., Ukleja, M., and Dziembowski, A. (2014). Proteins involved in the degradation of cytoplasmic mRNA in the major eukaryotic model systems. RNA Biol 11, 1122–1136.

Szczesny, R.J., Kowalska, K., Klosowska-Kosicka, K., Chlebowski, A., Owczarek, E.P., Warkocki, Z., Kulinski, T.M., Adamska, D., Affek, K., Jedroszkowiak, A., et al. (2018). Versatile approach for functional analysis of human proteins and efficient stable cell line generation using FLP-mediated recombination system. PloS one 13, e0194887.

Takagaki, Y., and Manley, J.L. (1998). Levels of polyadenylation factor CstF-64 control IgM heavy chain mRNA accumulation and other events associated with B cell differentiation. Molecular cell 2, 761–771.

Tang, A., C.S., van Baren M., Hart K., Hrabeta-Robinson E., Wu C., Brooks A. (2018). Full-length transcript characterization of SF3B1 mutation in chronic lymphocytic leukemia reveals downregulation of retained introns. BiorxiV, https://doi.org/10.1101/410183.

Tellier, J., Shi, W., Minnich, M., Liao, Y., Crawford, S., Smyth, G.K., Kallies, A., Busslinger, M., and Nutt, S.L. (2016). Blimp-1 controls plasma cell function through the regulation of immunoglobulin secretion and the unfolded protein response. Nat Immunol 17, 323–330.

Thai, T.H., Calado, D.P., Casola, S., Ansel, K.M., Xiao, C., Xue, Y., Murphy, A., Frendewey, D., Valenzuela, D., Kutok, J.L., et al. (2007). Regulation of the germinal center response by microRNA-155. Science 316, 604–608.

Tsuru A., Imai Y., Saito M., and Kohno K. (2016). Novel mechanism of enhancing IRE1α-XBP1 signalling via the PERK-ATF4 pathway. Scientific Reports 6, 24217

Vigorito, E., Perks, K.L., Abreu-Goodger, C., Bunting, S., Xiang, Z., Kohlhaas, S., Das, P.P., Miska, E.A., Rodriguez, A., Bradley, A., et al. (2007). microRNA-155 regulates the generation of immunoglobulin class-switched plasma cells. Immunity 27, 847–859.

Warkocki, Z., Liudkovska, V., Gewartowska, O., Mroczek, S., and Dziembowski, A. (2018). Terminal nucleotidyl transferases (TENTs) in mammalian RNA metabolism. Philos Trans R Soc Lond B Biol Sci 373.

Welch, J.D., Slevin, M.K., Tatomer, D.C., Duronio, R.J., Prins, J.F., and Marzluff, W.F. (2015). EnD-Seq and AppEnD: sequencing 3’ ends to identify nontemplated tails and degradation intermediates. Rna 21, 1375–1389.

Wiest, D.L., Burkhardt, J.K., Hester, S., Hortsch, M., Meyer, D.I., and Argon, Y. (1990). Membrane biogenesis during B cell differentiation: most endoplasmic reticulum proteins are expressed coordinately. The Journal of cell biology 110, 1501–1511.

Woo, Y.M., Kwak, Y., Namkoong, S., Kristjansdottir, K., Lee, S.H., Lee, J.H., and Kwak, H. (2018). TED-Seq Identifies the Dynamics of Poly(A) Length during ER Stress. Cell reports 24, 3630–3641 e3637.

Workman, R.E., Tang, A., Tang, P.S., Jain, M., Tyson, J.R., Zuzarte, P.C., Gilpatrick, T., Razaghi, R., Quick, J., Sadowski, N., et al. (2018). Nanopore native RNA sequencing of a human poly(A) transcriptome. BiorxiV, https://doi.org/10.1101/459529.

Xu, S., Guo, K., Zeng, Q., Huo, J., and Lam, K.P. (2012). The RNase III enzyme Dicer is essential for germinal center B-cell formation. Blood 119, 767–776.

Zhu, Y.X., Shi, C.X., Bruins, L.A., Jedlowski, P., Wang, X., Kortum, K.M., Luo, M., Ahmann, J.M., Braggio, E., and Stewart, A.K. (2017). Loss of FAM46C Promotes Cell Survival in Myeloma. Cancer Res 77, 4317–4327.

