## Supplementary_Tables for "B cell humoral response and differentiation is regulated by the non-canonical poly(A) polymerase TENT5C"

Supplementary table 1. Primer used for generation of TENT5C-TEV-GFP *knock-in* mice and genotyping.

| **Primer name** | **Sequence** |
| --- | --- |
| mTENT5C_GFP_gRNA_F | gaaattaatacgactcactatagggAGGTCTTCAGGTTAGTTACgttttagagctagaaatagcaagttaaaataaggc |
| Universal_gRNA_rev | CTTCAGAACCACTTCTCGGA |
| mTent5C_TOPO-LF_1f | CCACTAGTAACGGCCGCCAGTGTGCTGGAATTCtcgaGGAGATAACCCTGAAGGACA |
| mTent5C_LF-TEV_1r | accatgatatcaccctgaaaatacaaattctcGTTACAGGGCAGCCATG |
| mTent5C_eGFP-RF_1f | atcactctcggcatggacgagctgtacaagTAAcctgaagacctgaggg |
| mTent5C_RF-TOPO_1r | CGGCCGCCAGTGTGATGGATATCTGCAGAATTCttttcatactgggagtgacg |
| TEV_1f | gagaatttgtattttcagggtga |
| mCherry_eGFP_1r | TTActtgtacagctcgtcca |
| mTent5C_GFP_short_1F | TACATTGCGCACCCTCCAATTACC |
| mTent5C_GFP_short_1R | TACCTGAGAGCCCCTGCCCT |
| mTent5C_GFP_seq1F | CTTCAGAACCACTTCTCGGA |
| mTent5C_GFP_seq1R | AGAAGTCACGCCTCCTATTG |

Supplementary table 2. List of primers used in this study.

| Primer | Sequence | Species | Reference | Application |
| --- | --- | --- | --- | --- |
| Ig Kappa F | GTGCCTCAGTCGTGTGCTTC | Mouse | This study | qPCR |
| Ig Kappa R | TGCTGCTCATGCTGTAGGTG | Mouse | This study | qPCR |
| Ig Lambda F | AGACTTCGCCATCAGTCACC | Mouse | This study | qPCR |
| Ig Lambda R | CCAGTCCACTGTCACCACAC | Mouse | This study | qPCR |
| Ighm F | AACAGAGATCTGCATGTGCC | Mouse | This study | qPCR |
| Ighm R | TTCGTGGCCTCGCAGATGAG | Mouse | This study | qPCR |
| J chain F | GACGATGGTGTTCCTGAGAC | Mouse | This study | qPCR |
| J chain R | CAAGCTAGTCAGGGTAGCAAG | Mouse | This study | qPCR |
| GAPDH F | AAGGGCTCATGACCACAGTC | Mouse | [Kakiuchi-Kiyota](https://www.ncbi.nlm.nih.gov/pubmed/?term=Kakiuchi-Kiyota%20S%5BAuthor%5D&cauthor=true&cauthor_uid=21937740) *et al*., 2011 | qPCR |
| GAPDH R | GGATGACCTTGCCCACAG | Mouse | [Kakiuchi-Kiyota](https://www.ncbi.nlm.nih.gov/pubmed/?term=Kakiuchi-Kiyota%20S%5BAuthor%5D&cauthor=true&cauthor_uid=21937740) *et al*., 2011 | qPCR |
| TENT5C F | CAGTCACCTCCTCTTCCAACG | Mouse | This study | qPCR |
| TENT5C R | AACCTGATCCCAGTTGAGCAC | Mouse | This study | qPCR |
| PABPC1 F | TGCAGAGGATGGCAAGTGTACG | Mouse | Chorghade *et al.,* 2017 | qPCR |
| PABPC1 R | GCTAGGAGGATAGTATGCAGC | Mouse | Chorghade *et al.,* 2017 | qPCR |
| Xbp1 T F | TGGCCGGGTCTGCTGAGTCCG | Mouse | Oslowski i Urano, 2011 | qPCR |
| Xbp1 R1 | GTGTCAGAGTCCATGGGA | Mouse | This study | qPCR |
| Xbp1 S F | CTGAGTCCGAATCAGGTGCAG | Mouse | Oslowski i Urano, 2011 | qPCR |
| Xbp1 US F | CAGCACTCAGACTATGTGCA | Mouse | Oslowski i Urano, 2011 | qPCR |
| Xbp1 R2 | GTCCAACTTGTCCAGAATGCC | Mouse | This study | qPCR |
| PERK F | TGTCTTGGTTGGGTCTGATG | Mouse | This study | qPCR |
| PERK R | ACCGTTATCGTATGGATACTGG | Mouse | This study | qPCR |
| CHOP F | CTGCCTTTCACCTTGGAGAC | Mouse | Serrano *et al*., 2014 | qPCR |
| CHOP R | CGTTTCCTGGGGATGAGAT | Mouse | Serrano *et al*., 2014 | qPCR |
| GRP94 F | TCAAATCGAACACGGCTTGC | Mouse | This study | qPCR |
| GRP94 R | CCATGAAGTAGATTTTGTCC | Mouse | This study | qPCR |
| Ero1lB F | AAGTACTCGCAAGCAGCAAACAGC | Mouse | Aragon *et al*., 2012 | qPCR |
| Ero1lB R | TATCTCGCCCAGTCAATGAACGCT | Mouse | Aragon *et al*., 2012 | qPCR |
| Ire1 F | GCCGAAGTTCAGATGGAATC | Mouse | Tsuru et al., 2016 | qPCR |
| Ire1 R | ATCAGCAAAGGCCGATGA | Mouse | Tsuru et al., 2016 | qPCR |
| 5S rRNA F | CATACCACCCTGAACGCG | Human | This study | qPCR |
| 5S rRNA R | CTACAGCACCCGGTATTCCC | Human | This study | qPCR |
| 18S rRNA F | GAGAAACGGCTACCACATCCAA | Human | Kolesnikowa et al., 2018 | qPCR |
| 18S rRNA R | CCAATTACAGGGCCTCGAAAGA | Human | Kolesnikowa et al., 2018 | qPCR |
| 5.8S rRNA F | GGTGGATCACTCGGCTCGT | Human | Kolesnikowa et al., 2018 | qPCR |
| 5.8S rRNA R | CCGCAAGTGCGTTCGAAGTG | Human | Kolesnikowa et al., 2018 | qPCR |
| 28S rRNA F | GGGTGGTAAACTCCATCTAAGG | Human | Kolesnikowa et al., 2018 | qPCR |
| 28S rRNA R | GCCCTCTTGAACTCTCTCTTC | Human | Kolesnikowa et al., 2018 | qPCR |
| GAPDH F | ATCAAGAAGGTGGTGAAGCA | Human | Kobyłecki *et al*., 2018 | qPCR |
| GAPDH R | CATACCAGGAAATGAGCTTG | Human | Kobyłecki *et al*., 2018 | qPCR |
| ActB F | GCATGGGTCAGAAGGATTCC | Human | Eaton *et al*., 2018 | qPCR |
| ActB R | CCACACGCAGCTCATTGTAG | Human | Eaton *et al*., 2018 | qPCR |
| Ig Kappa F | CTGTCAGTCTTGGAGATCAAG | Mouse | This study | Northern blot probe |
| Ig Kappa R | CCGAACGTGTACGGAACATG | Mouse | This study | Northern blot probe |
| Ig Lambda F | GCTGTTGTGACTCAGGAATC | Mouse | This study | Northern blot probe |
| Ig Lambda R | GCTGACCTAGGACAGTGACC | Mouse | This study | Northern blot probe |

Supplementary table 3. List of pathogens detected in TENT5 KO and TENT5 WT mice.

| **Pathogen:** | **Method of detection:** | **Results 2018** | **Results 2017** |
| --- | --- | --- | --- |
| Ectromelia | Serology | Negative | Negative |
| EDIM | Serology | Weak positive | Negative |
| LCMV | Serology | Negative | Negative |
| *Mycoplasma pulmonis* | Serology | Negative | Negative |
| MAV1 | Serology | Negative | Negative |
| MAV2 | Serology | Negative | Negative |
| MHV | Serology | Positive | Positive |
| MNV | Serology | Positive | Positive |
| MPV | Serology | Negative | Negative |
| MVM | Serology | Negative | Negative |
| PVM | Serology | Negative | Negative |
| REO3 | Serology | Negative | Negative |
| TMEV | Serology | Negative | Negative |
| *Sendai* | Serology | Negative | Negative |
| *Clostridium piliforme* | PCR evaluation from feces | Negative | Negative |
| *Citrobacter rodentium* | PCR evaluation from feces | Negative | Negative |
| *Corynebacterium kutscheri* | PCR evaluation from feces | Negative | Negative |
| *Cryptosporidium spp.* | PCR evaluation from feces | Negative | Negative |
| *Helicobacter spp.* | PCR evaluation from feces | Positive | Positive |
| *Helicobacter bilis* | PCR evaluation from feces | Negative | Negative |
| *Helicobacter ganmani* | PCR evaluation from feces | Positive | Positive |
| *Helicobacter hepaticus* | PCR evaluation from feces | Negative | Negative |
| *Helicobacter mastomyrinus* | PCR evaluation from feces | Negative | Negative |
| *Helicobacter rodentium* | PCR evaluation from feces | Negative | Negative |
| *Helicobacter typhlonius* | PCR evaluation from feces | Positive | Positive |
| *Pasteurella pneumotropica biotype Jawetz* | PCR evaluation from feces | Positive | Positive |
| *Pasteurella pneumotropica biotype Heyl* | PCR evaluation from feces | Positive | Positive |
| *Salmonella spp.* | PCR evaluation from feces | Negative | Negative |
| *Streptobacillus moniliformis* | PCR evaluation from feces | Negative | Negative |
| *Streptococcus pneumoniae* | PCR evaluation from feces | Negative | Negative |
| *Streptococcus sp. beta hemolytic Group A* | PCR evaluation from feces | Negative | Negative |
| *Streptococcus sp. beta hemolytic Group B* | PCR evaluation from feces | Negative | Negative |
| *Streptococcus sp. beta hemolytic Group C* | PCR evaluation from feces | Negative | Negative |
| *Streptococcus sp. beta hemolytic Group G* | PCR evaluation from feces | Negative | Negative |
| *Giardia muris* | PCR evaluation from feces | Negative | Negative |
| *Spironucleus muris* | PCR evaluation from feces | Positive | Positive |
| *Aspiculuris tetraptera* | PCR evaluation from feces | Positive | Positive |
| *Syphacia muris* | PCR evaluation from feces | Negative | Negative |
| *Syphacia obvelata* | PCR evaluation from feces | Positive | Positive |
| *Myocoptes* | Pelt swap | Positive | Positive |
| *Radfordia/Myobia* | Pelt swap | Positive | Positive |
